# Therapy-induced senescent-like cancer cells drive macrophage-mediated immunosuppression in cholangiocarcinoma

**DOI:** 10.64898/2026.06.24.734341

**Authors:** Binbin Li, Jingchun Yang, Meina Cai, Shelby K. Yee, Danielle M. Carlson, Rory L. Smoot, Darren J. Baker, Sumera I. Ilyas

**Author notes:** Corresponding author: Sumera I. Ilyas, Associate Professor of Medicine, Mayo Clinic College of Medicine and Science, 200 First Street SW, Rochester, MN 55905.

## Abstract

Cholangiocarcinoma (CCA) is a lethal biliary cancer in which chemoresistance is nearly universal, and its tumor immune microenvironment is dominated by immunosuppressive tumor-associated macrophages (TAMs) that exclude cytotoxic CD8^+^ T cells. How tumor cells sustain this immunosuppressive state during chemotherapy is undefined. Here we demonstrate that therapy-induced senescent-like (Sen-L) cancer cells accumulate after gemcitabine/cisplatin in human and murine CCA, are predominantly cancer cells, and predict shorter survival. Genetic elimination of Sen-L cancer cells reduces tumor burden, lowers TAM abundance, and restores intratumoral CD8^+^ T cells, establishing them as causal drivers. Growth differentiation factor 15 (GDF-15) is the dominant Sen-L-secreted factor and reprograms macrophages to suppress CD8^+^ T cells through the non-canonical receptor TGFBR2 and STAT6, and p16-restricted *Gdf15* silencing phenocopies Sen-L elimination. Combined with chemotherapy, Sen-L elimination improves survival beyond chemotherapy alone. These findings establish Sen-L cancer cells and their GDF-15 output as causal, targetable drivers of macrophage-mediated immune evasion in CCA.

**SIGNIFICANCE STATEMENT:** Therapy-induced senescent-like cancer cells, not stromal cells, are the dominant senescent-like and immunosuppressive population in cholangiocarcinoma, and their elimination restores antitumor immunity. GDF-15 is their dominant secreted effector and engages a non-canonical macrophage receptor, TGFBR2, identifying a cancer-cell-to-macrophage axis and a Sen-L-elimination strategy to restore chemosensitivity.

**Graphical abstract.** Senescent-like CCA cells promote tumor immunosuppression through TAMs polarization by GDF-15

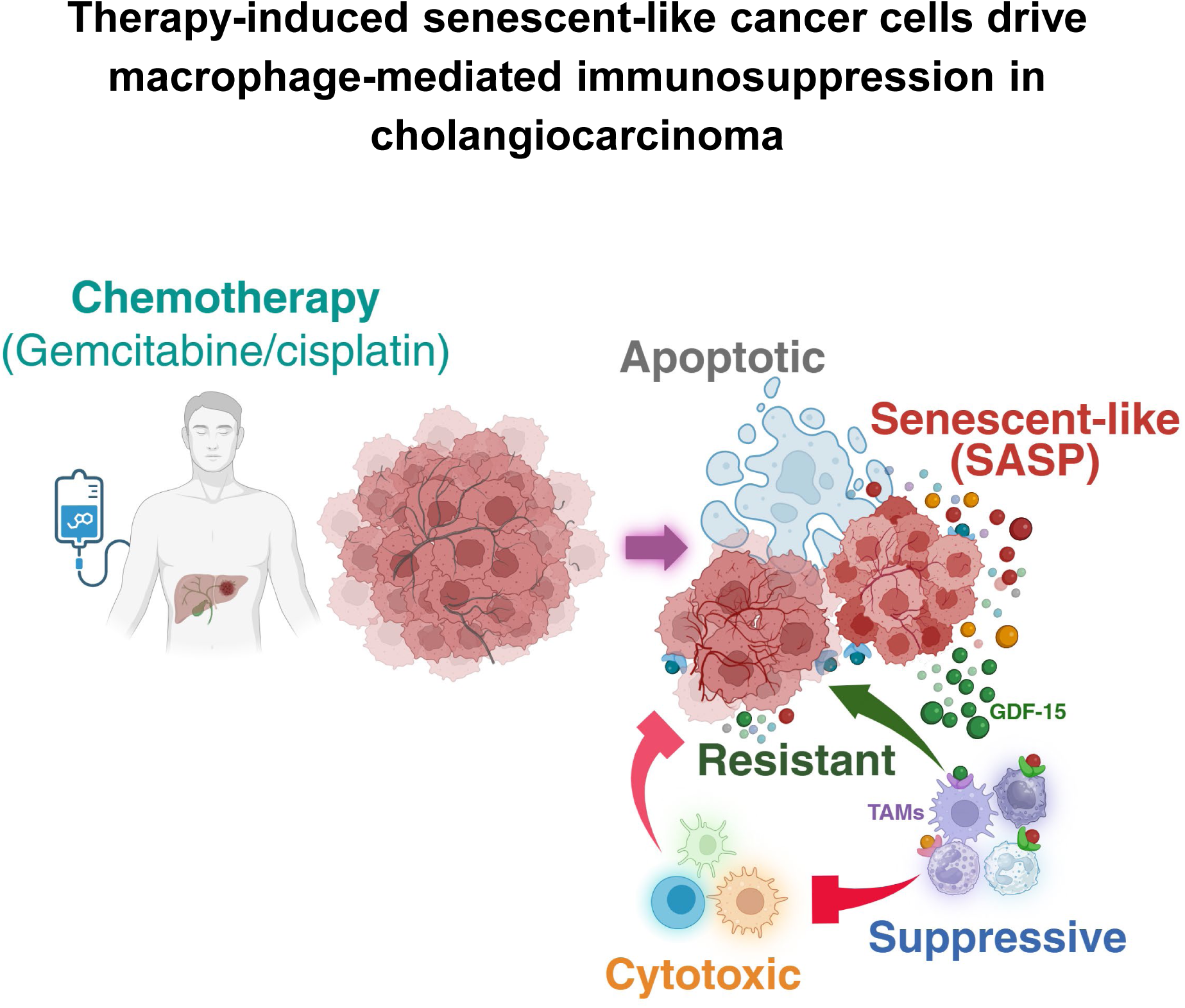

## INTRODUCTION

Cholangiocarcinoma (CCA) is a lethal biliary epithelial cancer with rising incidence and a five-year survival below 15%.^1^ For advanced disease, first-line gemcitabine plus cisplatin (Gem/Cis) combined with immune checkpoint inhibition yields a median overall survival of only about 12.8 months, and most patients progress on treatment.^2–3^ The addition of programmed death-ligand 1 (PD-L1) blockade to chemotherapy has produced only incremental survival gains, underscoring how poorly advanced CCA responds to current cytotoxic and immunotherapeutic regimens. Chemoresistance is the dominant cause of treatment failure, and durable benefit will require strategies that directly target the cellular drivers of resistance rather than the bulk tumor alone.^4^

Chemotherapy does not eliminate all malignant cells; it spares drug-tolerant cells that enter therapy-induced senescence, an established but therapeutically untargeted resistance mechanism.^5–9^ Senescent cells arise in tumors after genotoxic and mitotic stress, persist through a stable proliferative arrest, and can both limit the efficacy of cytotoxic therapy and establish a microenvironment that promotes immunotherapy resistance.^5–9^ Despite their recognized contribution to relapse, their role as causal drivers of tumor progression, as opposed to passive bystanders of treatment, has remained poorly defined, in part because senescent cells are difficult to identify unambiguously and have historically been studied as a tumor-suppressive endpoint of oncogene activation.

The consequences of senescence for antitumor immunity are context-dependent. In some experimental settings, senescent cells are immunogenic and promote antitumor immunity through enhanced antigen presentation and interferon-γ sensing, particularly when senescence is acute or vaccination-delivered in surveillance-permissive contexts.^10–11^ Through the senescence-associated secretory phenotype (SASP), however, persistent senescent cells remodel their surroundings, and many SASP factors are tumor-promoting.^12^ The net effect of senescence therefore varies with SASP composition, the chronicity of the senescent state, and the cellular compartment in which senescence occurs, reconciling the apparently opposing roles reported across tumor types.^13^ Which outcome dominates in CCA, a chemoresistant cancer treated with senescence-inducing agents, has not been established.

The CCA tumor immune microenvironment (TiME) is a poorly immunogenic but cellularly rich compartment sustained by tumor-associated macrophages (TAMs) that exclude cytotoxic T lymphocytes (CTLs) and enforce therapeutic resistance.^14–17^ The cancer-cell-derived signals that drive TAM reprogramming in CCA remain poorly defined. Senescent stromal cells, including fibroblasts and myeloid cells, drive immunosuppression in several solid tumors, among them pancreatic, breast, and prostate cancer and the senescent-macrophage compartment of the lung.^18–22^ The role of senescent-like cancer cells, as opposed to senescent stroma, in reprogramming the CCA TiME and enforcing chemoresistance is unknown, leaving open the possibility that the malignant epithelial compartment is itself the dominant source of immunosuppressive senescence.

Growth differentiation factor 15 (GDF-15) is a stress-induced TGF-β superfamily cytokine with a well-documented protumor role in cancer^23–24^ and a component of the SASP in aging.^25–26^ GDF-15 drives tumor immune evasion across cancer types and is now targeted by neutralizing agents in clinical trials.^24,27^ Its only established high-affinity receptor, GFRAL, is restricted to hindbrain neurons and is absent on immune cells, including tissue macrophages, on which GDF-15 nonetheless exerts clear functional effects.^28–29^ The receptor through which GDF-15 acts on macrophages has not been biochemically identified, although inflammatory-disease studies link GDF-15 to macrophage function^30^ and implicate the type II TGF-β receptor TGFBR2 in GDF-15 responses.^31^ Identifying the macrophage receptor for GDF-15 therefore carries significance well beyond CCA, given the advanced clinical development of GDF-15-directed therapeutics.

Herein, we demonstrate that therapy-induced senescent-like (Sen-L) cancer cells accumulate after chemotherapy in human and murine CCA, predict shorter overall survival, and are causal drivers of tumor progression and macrophage-mediated CD8^+^ T-cell suppression through GDF-15 engaging macrophage TGFBR2. Defining Sen-L cells functionally by their protumor secretome rather than by any single marker, we show that their selective genetic elimination reduces tumor burden, reverses TAM-mediated immunosuppression, and, combined with chemotherapy, improves survival. These findings reframe CCA chemoresistance from a cell-autonomous drug-tolerance problem to a non-cell-autonomous mechanism driven by a therapy-induced cancer-cell population, and they nominate the senescent-like cancer cell as a therapeutic target.

## RESULTS

### Sen-L cancer cells accumulate after chemotherapy and predict poor survival in CCA

Cytotoxic chemotherapy rarely eradicates CCA, prompting us to ask which cancer cells persist after treatment and how they reshape the immune microenvironment. By multiplex and sequential immunofluorescence on human CCA tumors, p16^+^ senescent-like (Sen-L) cancer cells were markedly more abundant in Gem/Cis-treated than in treatment-naive tumors (Fig. 1A and B), and a parallel rise in p21^+^ cancer cells (Supplementary Fig. S1A) indicated that chemotherapy induces this state across senescence markers. As an orthogonal hallmark of senescence, Gem/Cis also increased telomere-associated foci (TAF), persistent telomeric DNA-damage foci detected by combined γH2AX immunofluorescence and telomere fluorescence in situ hybridization, in CCA cancer cells (Fig. 1C).

**Figure 1.**
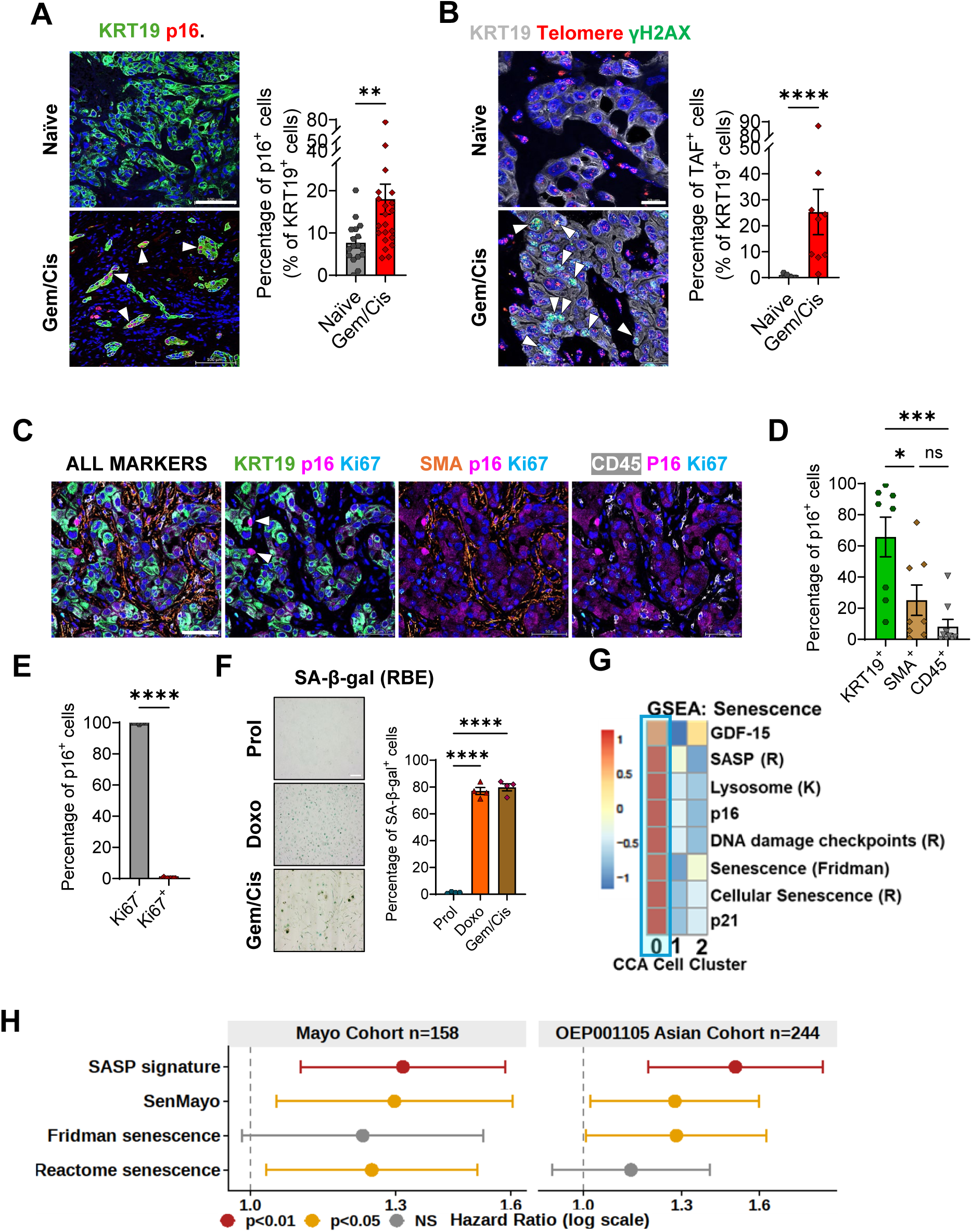
Senescent-like cancer cells accumulate after chemotherapy and predict poor survival in cholangiocarcinoma. (A) Representative images (left panel) and quantification (right panel) of p16^+^ cancer cells in treatment-naive (n=17) and gemcitabine/cisplatin (Gem/Cis)-treated (n=22) human CCA. (B) Telomere-associated foci (γH2AX immunofluorescence with telomere fluorescence in situ hybridization) (left panel) and quantification (right panel) in p16^+^ cancer cells. (C-E) Sequential immunofluorescence quantifying p16^+^ cancer cells (KRT19^+^), cancer-associated fibroblasts (α-SMA^+^), and immune cells (CD45^+^), and the proliferative status (Ki67) of p16^+^ cells. (F) Senescence-associated β-galactosidase activity in human RBE CCA cells, proliferating versus doxorubicin or Gem/Cis-induced. (G) Single-nucleus RNA sequencing of human CCA (n=10 patients): UMAP of cell populations and expression of *CDKN2A*, *CDKN1A*, and *MKI67* with senescence (SenMayo, Fridman, and Reactome cellular senescence) gene-set scores. (H) Multivariable Cox regression of senescence transcriptional scores and overall survival, adjusted for age, sex, and stage, performed independently in the Mayo discovery cohort (n=158, 108 events) and an external Asian cohort (OEP00105, n=244, 99 events). White arrows, senescent-like cancer cells. Scale bar, 50 µm. Data are mean ± SEM; *, P<0.05; **, P<0.01; ***, P<0.001; ****, P<0.0001; ns, not significant.

To determine which cells acquire these features, we localized the Sen-L compartment: the p16^+^ population was overwhelmingly composed of KRT19^+^ cancer cells rather than α-SMA^+^ cancer-associated fibroblasts or CD45^+^ immune cells and was almost uniformly non-proliferative (Ki67-negative) (Fig. 1C-E), with the same cancer-cell-dominant distribution for p21^+^ cells (Supplementary Fig. S1B-D). This cancer-cell-dominant pattern distinguishes CCA from the stromal-senescence paradigm of other solid tumors and focuses attention on the malignant epithelial compartment.

Critically, chemotherapy directly induced this Sen-L state in vitro. Gem/Cis treatment of human RBE CCA cells induced senescence-associated β-galactosidase (SA-β-gal) activity (Fig. 1F), and the induction was not idiosyncratic to one regimen or species, with doxorubicin, alisertib, and etoposide each eliciting the same activity across human and murine CCA cells (Supplementary Fig. S1E-F). The distinction between bona fide senescence and stress-induced states cannot be resolved in fixed tissue,^32^ so we define these Sen-L cancer cells by composite senescence criteria, including p16, growth arrest, telomere-associated foci, and senescence transcriptional signatures, together with their protumor secretome rather than by any single marker.^33^

Single-nucleus RNA sequencing of human CCA (n=10) corroborated this Sen-L state at single-cell resolution, resolving a discrete cancer-cell cluster enriched for senescence transcriptional programs (SenMayo,^34^ Fridman,^35^ and Reactome cellular senescence^36^) and depleted of *MKI67* (Fig. 1G). Clinically, Sen-L burden carried independent prognostic weight. Of four senescence transcriptional scores computed on bulk RNA-seq (SASP, SenMayo, Fridman, and Reactome), higher SASP and SenMayo scores were independently associated with shorter overall survival in multivariable Cox regression adjusting for age, sex, and AJCC stage, and were significant in both the Mayo discovery cohort and an independent Asian cohort (Fig. 1H). The reproducibility of this association across two geographically distinct cohorts, independent of stage, indicates that Sen-L burden is not merely a correlate of advanced disease. These data establish that therapy-induced Sen-L cells in CCA are predominantly cancer cells, arise across human and murine models in response to diverse chemotherapeutics, and independently identify patients with worse outcome.

### Genetic elimination of Sen-L cancer cells reduces CCA tumor burden

Having identified Sen-L cancer cells as a prognostic, chemotherapy-induced population, we next asked whether they are causally required for tumor progression or are merely a bystander of treatment. To distinguish these possibilities, we engineered a p16-promoter *INK-ATTAC* construct in the syngeneic murine SB1 CCA model,^37–38^ in which AP20187 (AP) drives dimerization of caspase-8 to selectively eliminate p16^+^ Sen-L cells while sparing transgene-negative stroma (Fig. 2A). *INK-ATTAC* reports execution of the senescence program rather than p16 expression per se,^39^ so proliferating p16^+^ cells are spared, providing specificity for the arrested Sen-L population. Sen-L cells constitute only ∼15% of cancer cells in SB tumors (Fig. 2B). In orthotopic SB1 *p16-ATTAC* tumors, AP markedly reduced tumor weight and volume relative to vehicle (Fig. 2C) (approximately 60% reduction), and elimination of p16^+^ Sen-L cells was confirmed by loss of the p16-promoter turbo green fluorescent protein (tGFP) reporter and reduced p16 mean fluorescence intensity (Fig. 2D and E).

**Figure 2.**
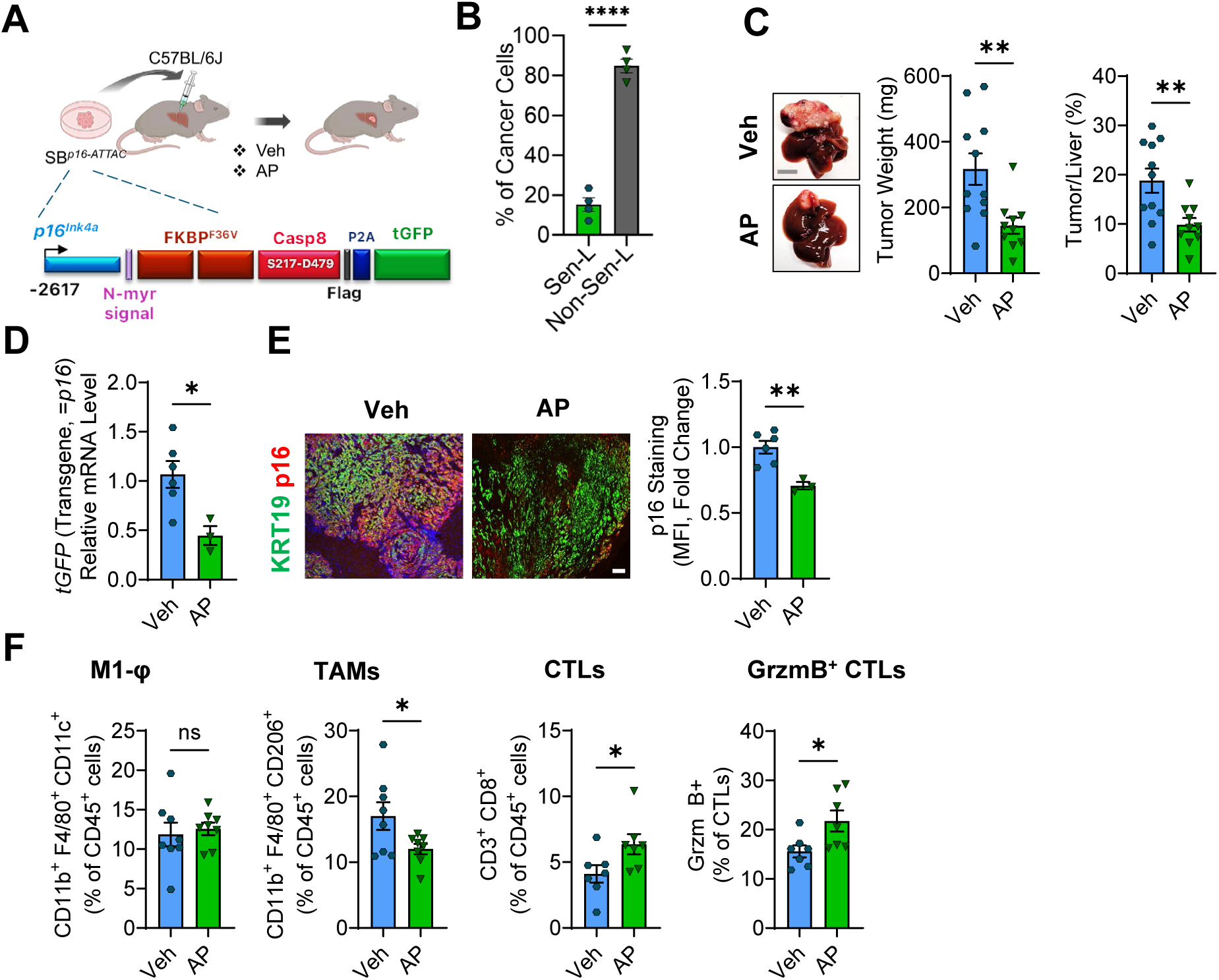
Genetic elimination of senescent-like cancer cells reduces cholangiocarcinoma tumor burden. (A) *p16-INK-ATTAC* strategy: AP20187 (AP) dimerizes caspase-8 to selectively eliminate p16^+^ senescent-like cancer cells while sparing transgene-negative stroma. (B) Fraction of cancer cells that are Sen-L (p16⁺ senescence-program⁺; ∼15%) in murine SB tumors (C) Representative gross images (left) and tumor-burden quantification (right) of orthotopic SB1 *p16-ATTAC* tumors treated with vehicle or AP (treatment from 2 weeks, harvest at 4 weeks post-implantation). (D) Relative mRNA of p16-promoter turbo green fluorescent protein (tGFP) reporter in veh and AP-treated murine SB1 *p16-ATTAC* tumors. (E) p16 mean fluorescence intensity in murine veh versus AP-treated SB1 *p16-ATTAC* tumors. (F) Frequencies of M1 macrophages, TAMs, CD8^+^ cytotoxic T lymphocytes (CTLs), and granzyme B^+^ CTLs in SB1 *p16-ATTAC* tumors. Scale bar, 100 µm. Data are mean ± SEM; *, P<0.05; **, P<0.01.

The magnitude of the tumor-burden reduction was disproportionate to the size of the targeted population: ablating only approximately 15% of cancer cells produced a far larger loss of tumor burden, a relationship incompatible with a purely cell-autonomous contribution and pointing instead to a non-cell-autonomous mechanism in which Sen-L cells act on the surrounding tumor. An independent p21-promoter *ATTAC* construct yielded convergent elimination and tumor reduction (Supplementary Fig. S2A-D), excluding an idiosyncrasy of the p16 reporter and demonstrating that the requirement tracks with the senescence program rather than with a single marker. Together, these data establish Sen-L cancer cells as causal drivers of CCA tumor burden.

### Sen-L cancer cells drive immunosuppressive TAM accumulation and CD8^+^ T-cell suppression

The disproportionate, non-cell-autonomous effect of Sen-L elimination implied that these cells act on the surrounding immune microenvironment rather than on tumor cells alone. We therefore profiled tumor immune composition by flow cytometry and tested macrophage reprogramming in vitro. Notably, the Sen-L-dependent immune remodeling was confined to the macrophage-CD8^+^ T cell axis. *INK-ATTAC* elimination of Sen-L cells reduced TAMs while restoring intratumoral CD8^+^ CTLs and granzyme B^+^ CTLs (Fig. 2F), linking the Sen-L population directly to both the suppressive myeloid compartment and the depth of cytotoxic infiltration. Conversely, Sen-L elimination in both *p16-ATTAC* and *p21-ATTAC* tumors left M-MDSC, PMN-MDSC, regulatory T-cell, CD4^+^ T-cell, and NK-cell frequencies unchanged (Fig. 2F, Supplementary Fig. S2E-H), arguing for a specific macrophage-directed program rather than a global shift in immune content. In orthotopic SB1 CCA (Fig. 3A), accumulation of Sen-L cells after chemotherapy was accompanied by an expansion of total macrophages and M2-like TAMs, without a comparable expansion of monocytic (M-MDSC) or polymorphonuclear (PMN-MDSC) myeloid-derived suppressor cells, or regulatory T cells (Fig. 3B and Supplementary Fig. S3A). The same increased-TAM, suppressed-CTL pattern recurred in a subcutaneous model (Supplementary Fig. S3B-C).

**Figure 3.**
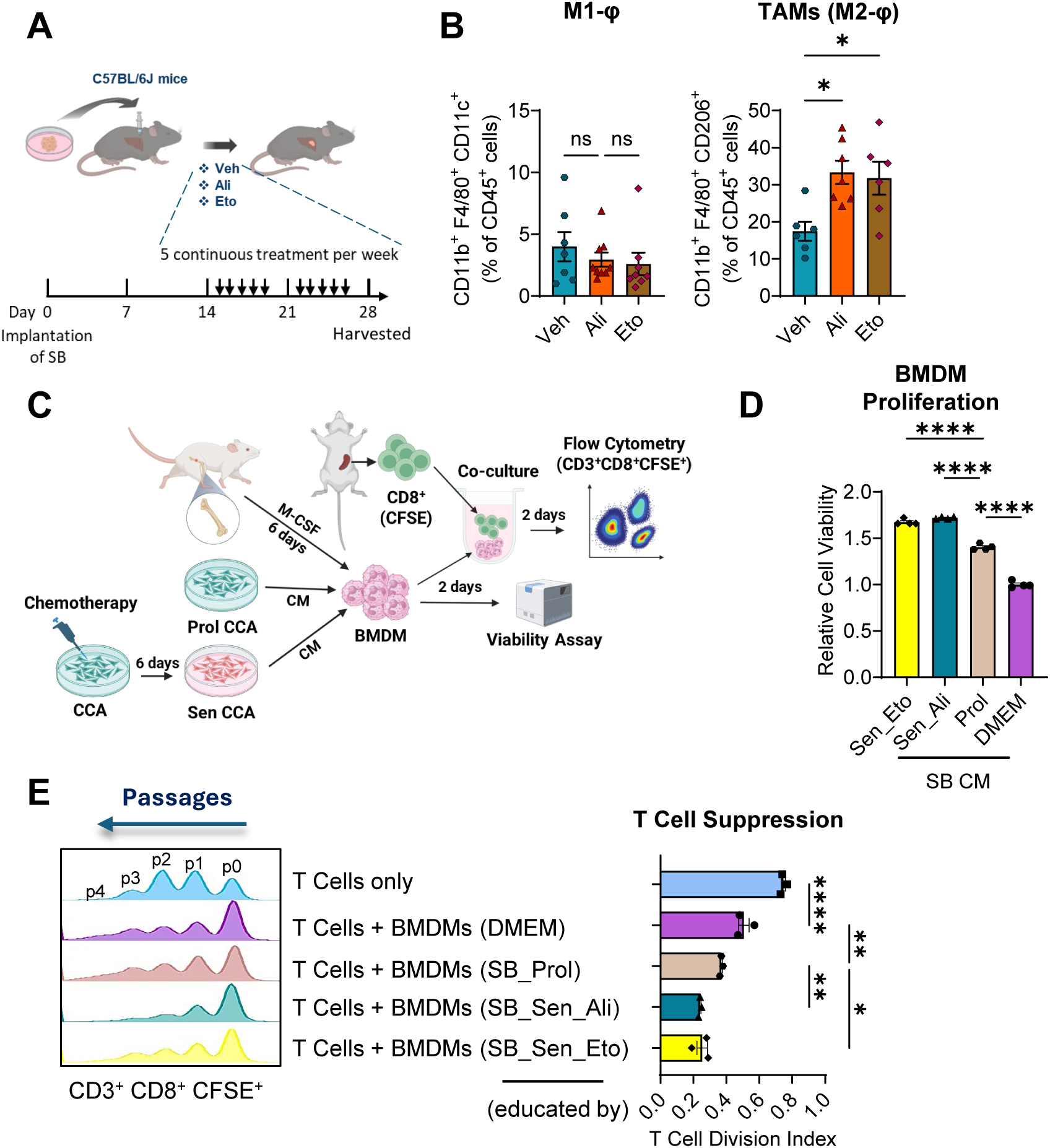
Senescent-like cancer cells drive immunosuppressive tumor-associated macrophage accumulation and CD8^+^ T-cell suppression. (A) Schema depicting in vivo study. SB tumor-bearing mice were treated for 2 weeks with vehicle, alisertib or etoposide. (B) Frequencies of M1 macrophages and M2-like tumor-associated macrophages (TAMs) (% of CD45^+^ cells) by flow cytometry. (C) Schematic of macrophage education; senescent-like state induced by alisertib or etoposide. (D) Bone-marrow-derived macrophage (BMDM) viability after conditioned medium from proliferating versus senescent-like cells. (E) Proliferation of CFSE-labeled CD8^+^ T cells co-cultured with educated BMDMs. Data are mean ± SEM; *, P<0.05; **, P<0.01; ***, P<0.001; ****, P<0.0001; ns, not significant.

To establish that Sen-L cancer cells reprogram macrophages directly rather than through other tumor-resident cells, we educated bone marrow-derived macrophages (BMDMs) with conditioned medium from proliferating or Sen-L CCA cells, the latter induced by alisertib or etoposide and confirmed by SA-β-gal and p21 staining (Fig. 3C). Sen-L-cell conditioned medium increased BMDM viability (Fig. 3D) and, after education, endowed BMDMs with the capacity to suppress carboxyfluorescein succinimidyl ester (CFSE)-labeled CD8^+^ T-cell proliferation (Fig. 3E), reconstituting the in vivo phenotype with a secreted signal alone. Collectively, these data demonstrate that Sen-L cancer cells reprogram macrophages into an immunosuppressive, CD8^+^ T cell-suppressive state across orthotopic and subcutaneous tumors and in vitro.

### Sen-L cancer-cell GDF-15 reprograms macrophages through TGFBR2 and STAT6

The non-cell-autonomous, macrophage-directed activity of Sen-L cells implicated a secreted mediator, prompting an unbiased survey of the Sen-L secretome. A 105-cytokine array of conditioned medium from proliferating versus doxorubicin- or Gem/Cis-induced senescent-like human RBE CCA cells identified GDF-15 as the most strongly induced factor (Fig. 4A). Consistent with a cancer-cell-autonomous source, *GDF15* and *Gdf15* transcripts were elevated in Sen-L human (RBE) and murine (SB1) CCA cells, respectively (Fig. 4B, Supplementary Fig. S4A), and *GDF15* was concentrated in the human single nucleus Sen-L cancer-cell cluster (Fig. 1G). Consistently, intratumoral *Gdf15* significantly decreased after Sen-L elimination in both ATTAC models (Fig. 4C, Supplementary Fig. S4B), tying the dominant secreted factor to the population whose removal reverses immunosuppression.

**Figure 4.**
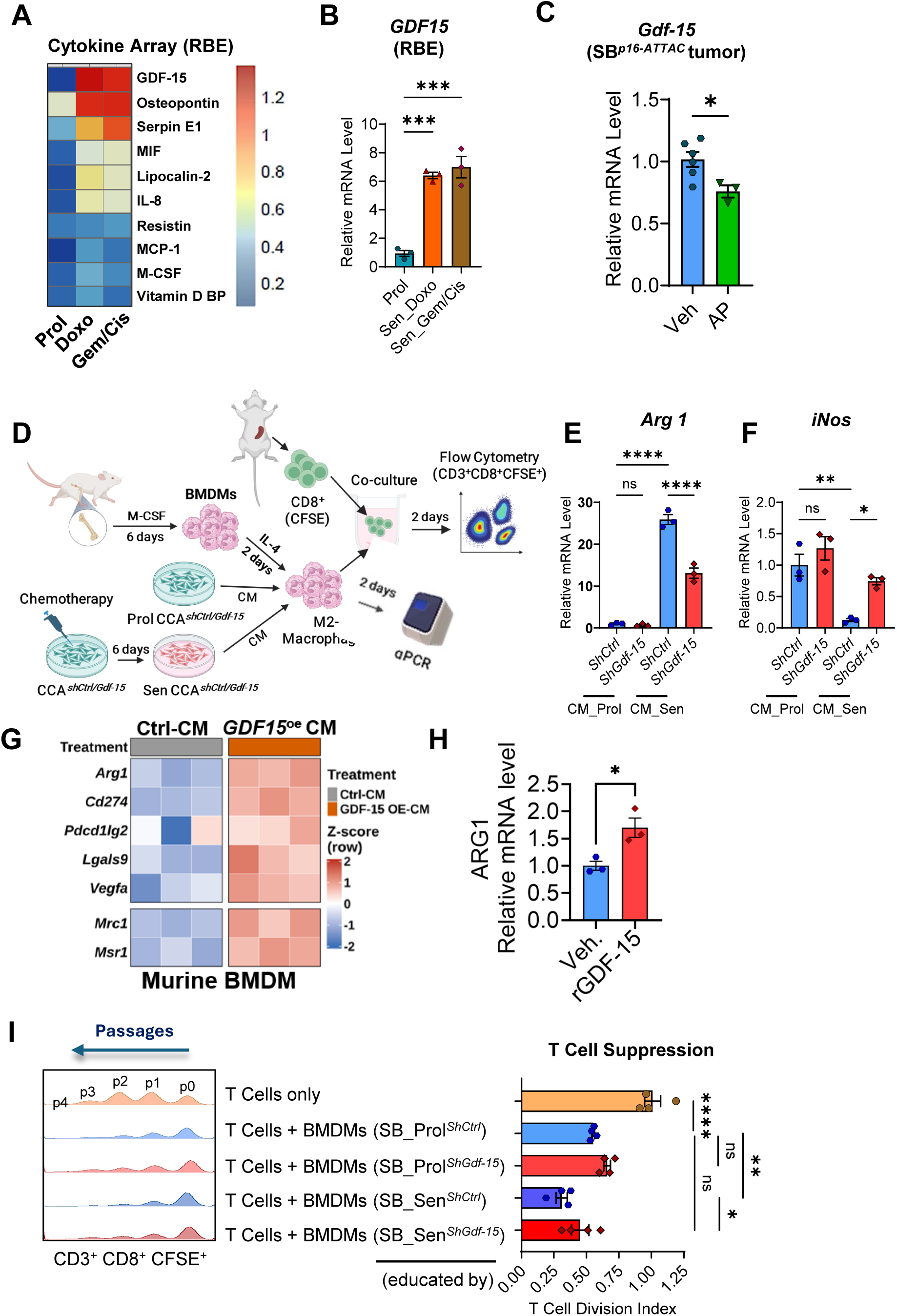
Senescent-cell-derived GDF-15 promotes macrophage immunosuppression. (A) Proteome Profiler Human XL cytokine array (105 cytokines) of conditioned medium from proliferating, doxorubicin-induced (Dox), and Gem/Cis-induced senescent-like RBE cells. (B) *Gdf15* mRNA in proliferating versus senescent-like murine CCA (SB1) cells. (C) *Gdf15* mRNA in control vs AP-treated *SB-p16-ATTAC* tumors. (D) Schematic of in vitro studies. (E-F) *Arg1* and *iNOS* mRNA in bone-marrow-derived macrophage (BMDM) incubated for 72 h with conditioned medium from vehicle- or alisertib-treated SB1 cells expressing *shCtrl* or *shGdf15*. (G) Bulk RNA-seq heatmap (row Z-scores) of naïve bone marrow-derived macrophages (BMDM) cultured 72 h with conditioned media (CM) from *SB-p16-shCtrl* or SB-*p16-Gdf15^OE^* cells. (H) *ARG1* mRNA in human monocyte-derived macrophages ± recombinant human GDF-15 (Qkine Qk017; TGF-β1-free), qRT-PCR relative to vehicle; mean ± SEM, n = 6. (I) Division index (CFSE dilution, normalized to T-cells-only) of anti-CD3/CD28-activated CD8⁺ T cells co-cultured with BMDMs incubated for 72 h with conditioned medium from vehicle- or alisertib-treated SB1 cells expressing *shCtrl* or *shGdf15*. Data are mean ± SEM; *, P<0.05; **, P<0.01; ****, P<0.0001; ns, not significant.

To test whether Sen-L-derived GDF-15 is functionally required, we silenced *Gdf15* with validated short hairpin RNA (shRNA) (Supplementary Fig. S4C); conditioned medium from *Gdf15*-silenced Sen-L cells no longer induced the macrophage immunosuppressive program (*Arg1*, *Spp1*), shifted macrophages toward an M1 profile (*iNOS*, *Cxcl9*), and relieved CD8^+^ T-cell suppression (Fig. 4D-F, Supplementary Fig. 4D-E). Bulk RNA sequencing demonstrated that conditioned medium from *Gdf15*-overexpressing murine cancer cells drove an immunosuppressive transcriptional program in naïve BMDMs (Fig. 4G). Recombinant human GDF-15 mediated immunosuppressive reprogramming in human CCA-patient monocyte-derived macrophages, as evidenced by increased *ARG1* expression (Fig. 4H), establishing direct GDF-15 sufficiency in human cells. Functionally, these Sen-L-CM-conditioned BMDMs suppressed CD8⁺ T-cell proliferation in co-culture, and *Gdf15* knockdown attenuated this suppression **(Fig. 4I)**. Together, these loss- and gain-of-function data establish cancer-cell GDF-15 as the dominant Sen-L-secreted factor and as functionally required for the macrophage reprogramming that suppresses CD8⁺ T cells.

We next asked how CCA macrophages sense GDF-15, whose only established receptor, GFRAL, is restricted to hindbrain neurons.^28–29^ Across mouse and human CCA single-cell datasets, macrophages lacked *GFRAL* (Fig. 5A, Supplementary Fig. S5A) but expressed *TGFBR2* (Fig. 5A-C). Proximity-dependent biotinylation from GDF-15 enriched TGFBR2 as a GDF-15-proximal macrophage receptor (Fig. 5C), identifying it as a candidate non-canonical receptor. Downstream, GDF-15 induced macrophage phospho-STAT6 without the canonical SMAD3 response (Fig. 5D), indicating that TGFBR2 signals non-canonically toward STAT6, the canonical activator of the immunosuppressive M2-like macrophage program. Functionally, *Tgfbr2* silencing in murine macrophages decreased the immunosuppressive marker *Arg1* (Fig. 5E, Supplementary Fig. S5B). Together, these data nominate macrophage TGFBR2 as a non-canonical receptor that transduces the GDF-15 signal to drive STAT6-dependent immunosuppression.

**Figure 5.**
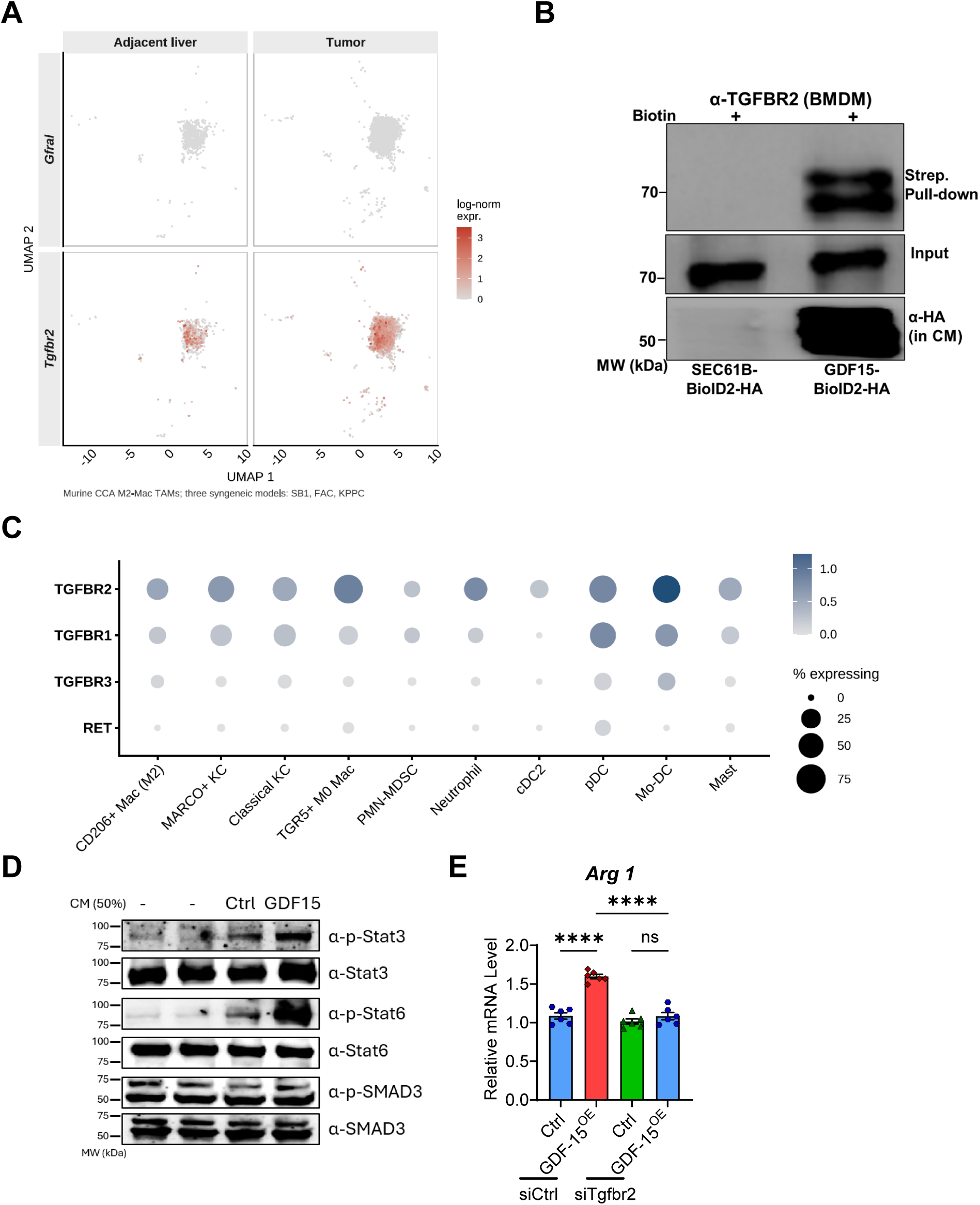
Senescent-like-cancer-cell GDF-15 reprograms macrophages through TGFBR2 and STAT6. (A) Macrophages express TGFBR2, the candidate non-canonical GDF-15 receptor, in murine and human CCA single-cell RNA-seq. (B) α-TGFBR2 western blot (WB) of streptavidin pull-down (top) and input (middle) from naïve BMDMs treated with biotin-supplemented conditioned media (CM) from CCA cells expressing GDF-15-BioID2-HA or SEC61B-BioID2-HA (ER control); α-HA blot of CM (bottom) confirms bait secretion. (C) GDF-15 induces macrophage phospho-STAT6 but not phospho-SMAD2/3, reflecting non-canonical TGFBR2 output. (D) WB of BMDMs stimulated with control or GDF-15-overexpressing (GDF-15^OE^) CM for 0 or 1 h: p-STAT6, STAT6, p-SMAD3, SMAD3. (E) *Arg1* mRNA in BMDMs transfected with *siCtrl* or *siTgfbr2* and stimulated with control or rGDF-15. n = 6. Data are mean ± SEM; *, P<0.05; **, P<0.01; ****, P<0.0001; ns, not significant.

### GDF-15 is required in vivo for TAM immunosuppression and CCA tumor growth

To test the in vivo requirement for Sen-L-derived GDF-15, we placed *Gdf15* silencing or overexpression under the p16 promoter in orthotopic SB1 CCA, restricting manipulation to the Sen-L compartment (Fig. 6A, Supplementary Fig. S6A). p16-restricted *Gdf15* silencing reduced tumor burden whereas *Gdf15* overexpression increased it, demonstrating that Sen-L-derived GDF-15 is both necessary and sufficient for tumor growth (Fig. 6B, Supplementary Fig. S6A). Mirroring the Sen-L genetic-elimination phenotype, *Gdf15* silencing lowered TAM abundance and restored intratumoral CD8^+^ CTLs (Fig. 6C-D), with concordant effects in a subcutaneous model (Supplementary Fig. 6C-D), indicating that the immunosuppressive output of Sen-L cells is conveyed largely through this single factor.

**Figure 6.**
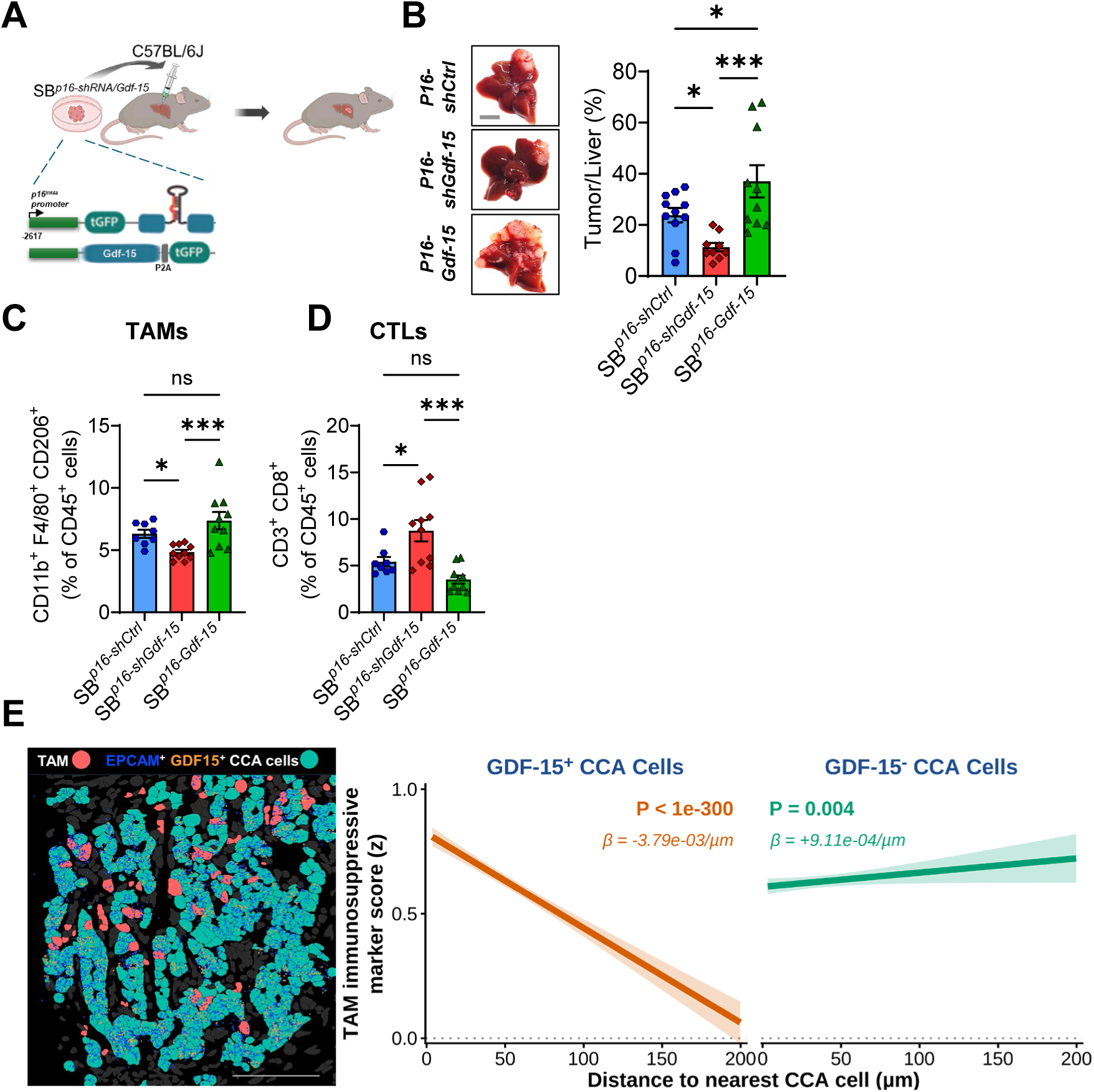
GDF-15 is required in vivo for tumor-associated macrophage immunosuppression and cholangiocarcinoma tumor growth. (A) Strategy for p16-promoter-restricted *Gdf15* silencing (shRNA) and *Gdf15* overexpression in orthotopic SB1 CCA. (B) Representative gross images (left) and tumor/liver weight quantification (right). (C-D) TAM and CD8^+^ CTL frequencies (% of CD45^+^ cells) in p16-shCtrl*, SB-p16-shGdf15, and* SB-*p16-Gdf15^OE^*tumors via flow cytometry. (E) (A) EpCAM⁺GDF-15⁺ CCA cells (teal) adjacent to TAMs (pink); EpCAM⁺ (blue) and GDF-15⁺ (orange) transcripts shown. Scale bar 100 µm. (B) Per-TAM immunosuppressive score (per-sample z) versus distance to the nearest GDF-15⁺ or GDF-15⁻ CCA cell (TAMs ≤200 µm; n = 13,597 and 14,703 across three iCCA samples). Line/band: linear fit ± 95% CI; β and P from linear mixed-effects [score ∼ distance + (1|sample)]. The gradient is GDF-15⁺-specific (β = −3.79×10⁻³/µm, P < 10⁻¹⁰) and absent for GDF-15⁻ cells (β = +9.11×10⁻⁴/µm, P = 0.004). Scale bar, 100 µm. Data are mean ± SEM; *, P<0.05; ***, P<0.001; ns, not significant.

To anchor the axis in human tissue, we examined its spatial organization by Xenium spatial transcriptomics. *GDF15*^+^ cancer cells localized adjacent to TAMs, and TAMs neighboring *GDF15*^+^ cancer cells exhibited higher immunosuppression scores than those near *GDF15*-negative cancer cells (Fig. 6E), placing the ligand-bearing cells and their macrophage targets in direct physical proximity; *GDF15*^+^ cancer-cell density likewise correlated with TAM proximity by GeoMx (Spearman ρ=0.63; Fig. 6D). Collectively, these data demonstrate that Sen-L-derived GDF-15 is necessary in vivo for TAM-mediated immunosuppression and CCA growth and that the cancer-cell-to-macrophage axis is spatially organized in human tumors.

### Sen-L elimination enhances chemotherapy

Finally, we asked whether targeting the Sen-L population is therapeutically actionable. In a head-to-head survival study, Gem/Cis plus genetic Sen-L elimination prolonged survival beyond Gem/Cis alone and beyond the current first-line standard of care, Gem/Cis plus anti-programmed cell death protein 1 (anti-PD-1) (Fig. 7A). Genetic elimination is not pharmacologically tractable, so we sought a druggable vulnerability of Sen-L cells. Reasoning that senescent cells survive through anti-apoptotic dependence, we tested BCL-2 family vulnerability: small interfering RNA (siRNA) against *Bcl2l1*, *Bcl2*, or *Mcl1*, validated by quantitative PCR (qPCR) (Fig. 7B), selectively reduced the viability of Sen-L but not proliferating cells (Fig. 7C),^40–41^ identifying a senescence-selective survival dependence. Together, these data identify Sen-L cancer cells as a BCL-2-family-dependent population whose elimination restores chemosensitivity and motivate tumor-restricted strategies to eliminate Sen-L cells.

**Figure 7.**
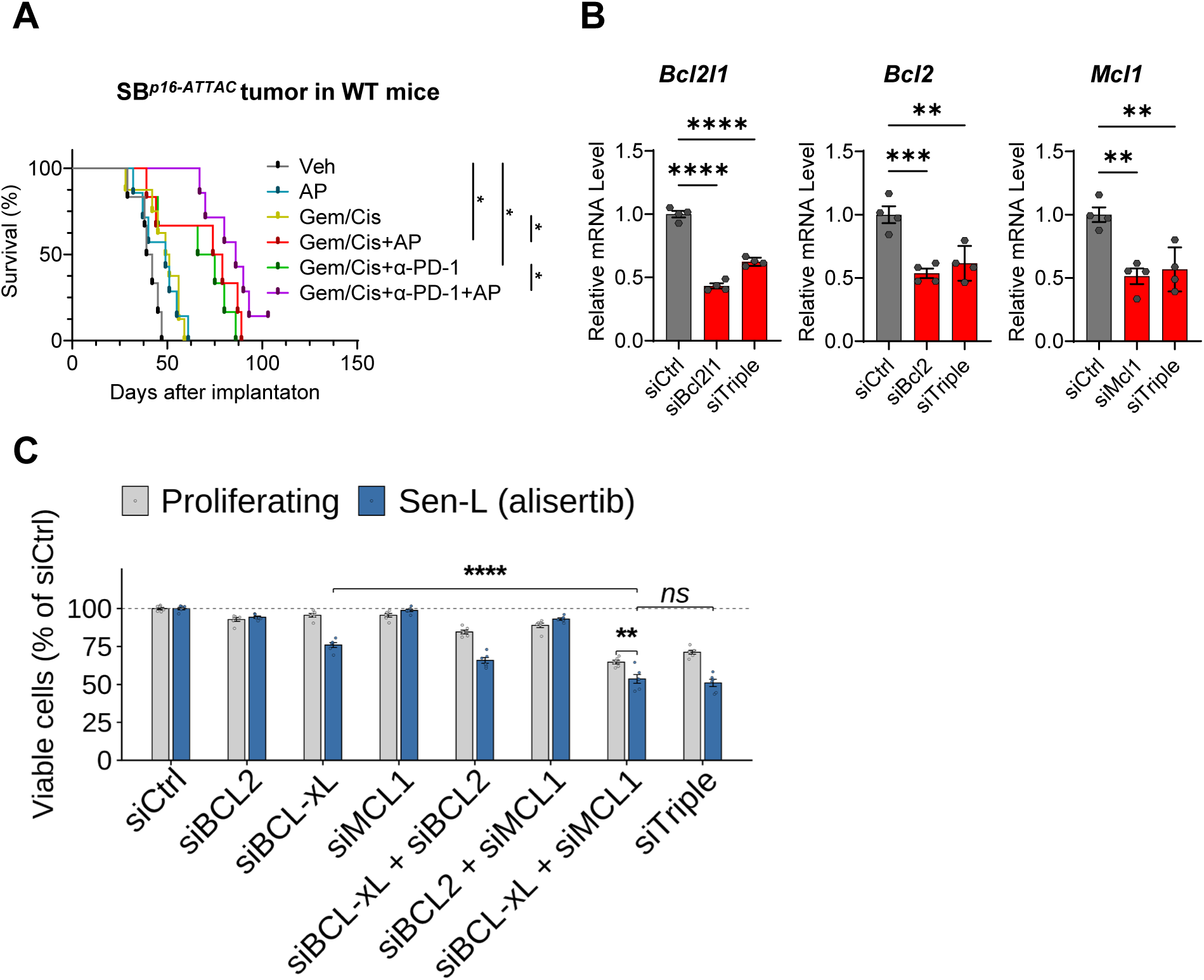
Sen-L elimination enhances efficacy of Gem/Cis chemotherapy. (A) *SB-INK-ATTAC* orthotopic tumor-bearing mice were treated with vehicle (Veh), AP, gemcitabine plus cisplatin (Gem/Cis), or Gem/Cis+AP. Mice were followed for survival. Kaplan-Meier curves show overall survival. Pairwise comparisons by log-rank (Mantel-Cox) test, *, P<0.05 (B) Viability of proliferating versus senescent-like SB1 cells after *Bcl2l1*, *Bcl2*, or *Mcl1* siRNA. (C) CCA cells transfected with the indicated siRNAs (siTriple = siBCL-xL + siBCL2 + siMCL1), treated with vehicle (proliferating) or alisertib (senescent); 48-h viability normalized to siCtrl proliferating. Data are mean ± SEM; *, P<0.05; **, P<0.01; ***, P<0.001, ****, P<0.0001; ns, not significant.

## DISCUSSION

We identify therapy-induced Sen-L cancer cells as causal drivers of macrophage-mediated immune evasion in cholangiocarcinoma. Sen-L cells are predominantly cancer cells, accumulate after chemotherapy, and independently predict worse survival; their genetic elimination reduces tumor burden, lowers TAM abundance, and restores intratumoral CD8^+^ T cells. Mechanistically, Sen-L cells act through GDF-15 engagement of macrophage TGFBR2 and downstream STAT6, and their selective elimination, combined with chemotherapy, improves survival beyond chemotherapy alone. These results reframe the immunosuppressive microenvironment that accompanies chemotherapy in CCA from a purely cell-autonomous drug-tolerance problem toward a non-cell-autonomous mechanism in which a therapy-induced cancer-cell population actively enforces treatment resistance.

Senescence has context-dependent roles in cancer.^13^ In some settings senescent cells are immunogenic and promote antitumor immunity through enhanced antigen presentation and interferon-γ sensing, particularly when senescence is acute or vaccination-delivered in surveillance-permissive contexts.^10–11^ In CCA we observe the opposite net outcome, in which chronic, therapy-induced Sen-L cancer cells persist in established tumors and elaborate an immunosuppressive, GDF-15-dominant SASP that acts on macrophages. This divergence is consistent with protumor senescent-cell programs reported in pancreatic, breast, and prostate cancer and in the senescent-macrophage compartment of the lung.^18–22^ Across these settings, the influence of senescence on antitumor immunity tracks with SASP composition, the chronicity of the senescent state, and the cellular compartment in which senescence arises rather than with the senescent state per se,^7–8,12–13^ and our place the chemotherapy-induced Sen-L state in CCA firmly within the immunosuppressive, tumor-promoting category.

A central finding is that, unlike the stromal-senescence paradigm in which senescent fibroblasts or myeloid cells orchestrate immunosuppression,^18–21^ the dominant senescent-like and immunosuppressive population in CCA is the malignant epithelial cell itself. Multiplex imaging localized the p16-high, non-proliferative compartment overwhelmingly to KRT19^+^ cancer cells rather than to stroma or immune cells, and single-nucleus profiling resolved a discrete senescence-high cancer-cell state conserved between human and murine tumors. The persistent difficulty of distinguishing bona fide senescence from other stress-induced states in fixed tissue has complicated efforts to assign function to senescent cancer cells.^32^ By defining Sen-L cells functionally, through the protumor secretome that reprograms the TiME rather than through any single marker, we circumvent this ambiguity and focus attention on the secreted output that is therapeutically actionable. This functional definition, anchored by the consequence of eliminating the population, distinguishes causal Sen-L cancer cells from incidental marker-positive cells.

The disproportionate reduction in tumor burden that followed elimination of a numerically small Sen-L population pointed to a non-cell-autonomous mechanism, which we traced to reprogramming of the macrophage compartment. Sen-L elimination selectively reduced immunosuppressive TAMs and restored cytotoxic CD8^+^ T cells while leaving MDSC, regulatory T-cell, CD4^+^ T-cell, and natural killer (NK)-cell frequencies unchanged, indicating a focused macrophage-CD8 axis rather than a global shift in immune content. That conditioned medium from Sen-L cells was sufficient to confer suppressive activity on BMDM establishes that the signal is secreted and acts directly on macrophages, independent of other tumor-resident cells. This positions the senescent cancer cell upstream of the TAM-enforced exclusion of CTLs that characterizes the CCA TiME and that we and others have linked to therapy resistance.^14–17^

GDF-15 drives tumor immune evasion across cancers and is being targeted clinically,^23–24,27^ yet the receptor through which it acts on macrophages has remained undefined because its canonical receptor GFRAL is restricted to the brainstem and absent on immune cells.^28–29^ We find that Sen-L-cancer-cell GDF-15 engages macrophage TGFBR2 and signals through STAT6, defining a non-canonical, cancer-cell-to-TAM ligand-receptor axis. This is supported by prior reports linking GDF-15 to macrophage function^30^ and to TGFBR2-dependent responses in non-tumor and other tumor contexts,^31,42–43^ and it is mechanistically distinct from the GDF-15/CD48 axis described on regulatory T cells in hepatocellular carcinoma.^44^ The phosphorylation of STAT6 rather than canonical SMAD2/3 effectors is consistent with the M2-like macrophage program that GDF-15 elicits and indicates that the receptor is repurposed toward a non-canonical output in this setting. With anti-GDF-15 agents already in clinical development,^24^ assigning a macrophage receptor to GDF-15 has implications beyond CCA, including the possibility that responses to GDF-15 blockade depend on TGFBR2 status in the myeloid compartment.

These observations support a model in which the immunosuppressive consequence of cancer-cell senescence is set jointly by the signal that Sen-L cells send and by how the recipient macrophage is equipped to receive it. On the sending side, the therapy-induced Sen-L program reshapes the cancer-cell secretome toward a GDF-15-dominant, immunosuppressive SASP. On the receiving side, the macrophage reads this signal not through the canonical GDF-15 receptor GFRAL, which it lacks, but through a repurposed TGFBR2 that redirects the output to STAT6 and an M2-like program. Cast as a coupled send-and-receive circuit, the axis predicts that immunosuppression requires both an appropriately composed Sen-L secretome and a macrophage poised to sense it, such that disrupting either arm, the senescent source or its non-canonical receptor, should interrupt the cancer-cell-to-TAM signal. This framework also accommodates the context-dependence of senescence in cancer, in which a secretory program can yield opposing immune outcomes depending on the receiving compartment and its sensing state.

Our findings argue for targeting the cellular source of GDF-15 rather than the ligand alone. Eliminating Sen-L cells removes the entire immunosuppressive secretome and forecloses compensation by other SASP factors, whereas neutralizing GDF-15 leaves the chemoresistant Sen-L reservoir intact and subject to escape through redundant mediators.^24^ Genetic, Sen-L-restricted elimination combined with chemotherapy improved survival while restoring intratumoral CD8^+^ T cells, and it exceeded the benefit of adding PD-1 blockade, indicating an immune-dependent rather than a purely cytoreductive gain. Senescent cells depend on BCL-2 family anti-apoptotic proteins for survival, a tractable vulnerability we confirmed in Sen-L CCA cells,^40–41^ yet systemic BCL-2 family inhibition carries dose-limiting, on-target thrombocytopenia and cardiotoxicity.^45–46^ Translating this dependence will therefore favor tumor- and Sen-L-directed delivery, for example a CCA-tropic nanoparticle carrying BCL-2-family-directed cargo, that concentrates the senolytic effect within the tumor while sparing systemic compartments. Such an approach remains to be tested and is a focus of ongoing work.

Beyond its role as a therapeutic target, Sen-L burden may also report on disease behavior. Sen-L transcriptional scores independently predicted shorter survival across two geographically distinct patient cohorts, and spatial transcriptomics placed *GDF15*^+^ cancer cells in direct physical proximity to the TAMs they reprogram, with neighboring macrophages displaying higher immunosuppression scores. The spatial organization of the axis in human tumors suggests that Sen-L density, measured in situ, could serve as a candidate biomarker of chemorefractory, immunosuppressed disease and could help identify the patients most likely to benefit from Sen-L-directed therapy. Prospective evaluation of Sen-L spatial burden against treatment response will be required to establish its clinical utility.

Several limitations frame these conclusions. First, senescence markers can label nonsenescent cells, and irreversible growth arrest cannot be established in fixed human tissue,^32^ so we define Sen-L cells by composite criteria, including p16, growth arrest, telomere-associated foci, and senescence transcriptional signatures,^33–36^ and adopt the conservative term senescent-like throughout. Second, the *INK-ATTAC* and *p21-ATTAC* systems eliminate cells executing the senescence program rather than all marker-positive cells per se,^38–39^ conferring specificity for the arrested population but not capturing senescent cells that have exited the program. Third, the macrophage GDF-15-TGFBR2 interaction is nominated here by proximity labeling and loss-of-function; its precise biochemical mode, stoichiometry, and in vivo genetic requirement remain to be defined. Finally, the therapeutic studies rely on genetic elimination, and translation will require a delivery strategy that achieves comparable Sen-L selectivity pharmacologically.

Together, these findings establish Sen-L cancer cells and their GDF-15 output as causal, targetable drivers of macrophage-mediated immune evasion in CCA. They nominate elimination of the Sen-L cancer-cell source as a candidate strategy to restore chemosensitivity and motivate further definition of the GDF-15-TGFBR2 axis and of Sen-L burden as an indicator of chemorefractory disease.

## METHODS

### Study Approval

Human CCA specimens were obtained from patients who provided written informed consent under Mayo Clinic Institutional Review Board (IRB) protocol 707-03; archival formalin-fixed, paraffin-embedded (FFPE) tumor tissue was studied under IRB 18-006218. All animal experiments were conducted under Mayo Clinic Institutional Animal Care and Use Committee (IACUC) oversight and adhered to the ARRIVE guidelines.^47^

### Mice

8-week-old male C57BL/6J mice (#000664) mice obtained from the Jackson Laboratory were used for all studies. Mice are housed under specific-pathogen-free conditions at Mayo Clinic with a stable temperature, humidity, and light cycle periods of 12 h in accordance with their circadian rhythm, and all animal procedures were reviewed and approved by the Mayo Clinic Institutional Animal Care and Use Committee.

### Murine CCA Model and treatment

Using a 27-gauge needle, 1 million SB1 cells were resuspended in 20 ul 1:1 mixture of unsupplemented DMEM and Matrigel (Corning #354263) and orthotopically implanted into the lateral lobe of the mouse liver under isoflurane anesthesia using aseptic technique, as previously described.^37^ The cotton tipped applicator was held over the injection site to prevent cell leakage and encourage hemostasis. Subsequently, the abdominal wall and skin were closed in separate layers with absorbable 4-0 Vicryl suture material and 3M™ Vetbond™ Tissue Adhesive, 1469SB. Tumor presence was confirmed using small animal ultrasound prior to treatment start. Mice were randomly and equally assigned to vehicle control or treatment groups. Treatment began two weeks after implantation, and tumors were harvested at four weeks. Gemcitabine (4 mg/kg) and cisplatin (1 mg/kg) were administered intraperitoneally twice weekly. Anti-PD-1 (10 mg/kg, clone G4.1 C9, Antibody Hybridoma Core, Mayo Clinic) were administered intraperitoneally every 3 days. AP20187 (10 mg/kg, MedChemExpress #HY-13992) were administered intraperitoneally every 3 days. For subcutaneous murine CCA models, 1 million SB1 cells were resuspended in 100 ul 1:1 mixture of unsupplemented DMEM and Matrigel were used, and alisertib (25 mg/kg) and etoposide (25 mg/kg) were administrated by oral gavage as indicated. Treatment groups were balanced for cage, surgical date, and passage number alongside age-matched vehicle controls, and starting tumor size was equalized across arms by Prospect T1 high-frequency ultrasound system (S-Sharp Corporation, New Taipei City, Taiwan)

### INK-ATTAC and p21-ATTAC Constructs

SB1 cells were stably engineered with an ATTAC cassette encoding an AP20187-inducible caspase-8 fused to a tGFP reporter, driven by a p16 (*INK-ATTAC*) or p21 promoter fragment, as previously described.^38–39^ AP20187 dimerizes caspase-8 to trigger apoptosis selectively in transgene-expressing Sen-L cells while sparing transgene-negative stroma; because the construct reports execution of the senescence program rather than p16 expression alone, proliferating p16-positive cells are spared.

### Gene Silencing and Overexpression

Cancer cell-derived GDF-15 was genetically manipulated using p16 promoter-driven constructs for senescence-dependent expression, and TRE3GS (inducible) or CMV promoter were used for constitutive expression. Stable knockdown was achieved using miR-E based shRNAs fused downstream of tGFP ORF targeting either *Renilla luciferase* as a control (shCtrl; guide sequence: 5′-TAGATAAGCATTATAATTCCTA-3′) or *Gdf-15* (shGdf15). Three independent shRNA sequences targeting *Gdf-15* were evaluated: #1, 5′-TAAGTCTGCAGTGACACACCAC-3′; #2, 5′-TTTAAATAAATACACAATCCAT-3′; and #3, 5′-TACAAATATAAATAGCATCCTT-3′. Unless otherwise indicated, shGdf15 #3 was used for all experiments. Mouse Gdf-15 ORF fused to tGFP was used for GDF-15 overexpression. Knockdown and overexpression efficiency were confirmed by quantitative RT-PCR in stably transfected SB cells.

For transient gene silencing, Horizon ON-TARGETplus non-targeting siRNA was used as a control. *Bcl2l1*, *Bcl2*, *Mcl1*, and *Tgfbr2* were silenced using gene-specific ON-TARGETplus siRNAs. Transfection was conducted using DharmaFECT 2 reagent in SB cells and DharmaFECT 1 reagent in BMDMs, according to the manufacturer’s instructions. Knockdown efficiency was verified by quantitative RT-PCR.

### Cell Culture

Human RBE CCA cells and the syngeneic murine CCA line SB1, derived from an oncogene-driven (YAP/Akt) biliary model as previously described,^37^ were maintained in high glucose DMEM supplemented with 10% fetal bovine serum and Primocin at 37°C in 5% CO₂. SB^p16/21-ATTAC^, SB^p16-shCtrl/shGdf-15/Gdf-15^ and SB^Ctrl/Gdf-15^ ^OE^ were generated by stable transfection with Sleeping Beauty transposase and respective pT3 based vectors with transposon elements carrying shRNA or gene cassette as described above, selected and maintained in high glucose DMEM supplemented with 10% fetal bovine serum and Primocin at 37°C in 5% CO₂.

Bone marrow derived macrophages (BMDMs) were isolated from the leg of 8-week non-tumor-bearing male C57BL/6J mice and maintained in Basal medium (RPMI 1640 with low endotoxin FBS and Primocin) supplemented with recombinant mouse M-CSF (20 ng/ml) for 6 days, and polarized into M1 or M2 phase by treating with LPS (20 ng/ml) or recombinant mouse IL4 (20 ng/ml) for another 2 days when needed.

CD8 T cells were isolated from splenocytes of 8-week male C57BL/6J mice and maintained in Basal medium supplemented with recombinant mouse IL2 (20 ng/ml) and activated by Dynabeads Mouse T-Activator CD3/CD28 (1:1).

### T-cell Suppression Assay

Conditioned medium (CM) from proliferating and senescent SB cells was collected by replacing the culture medium with serum-free DMEM and incubating the cells for 24 hours. CM samples were subsequently normalized to the final cell number at the time of collection to account for differences in cell density. BMDMs were generated as described above, and polarized when needed, and educated by indicated CMs for 48h without growth factors. BMDMs were then replated with CFSE (ThermoFisher # C34554) labeled CD8 T cells in basal medium with IL2 and CD3/28 beads at 1:4 ratio in 96 well “U” bottom plate for 3 days co-culture. Cells were harvested and stained with Fixable Viability Stain 660, anti-CD3, and anti-CD8 antibodies as described in the “Flow Cytometry” section. T-cell proliferation was quantified by determining the division index in FlowJo software.

### Senescent-like State Induction

The senescent-like (Sen-L) state was induced in CCA cells with gemcitabine (10 nM) plus cisplatin (7 µM), doxorubicin (0.5 µM), the Aurora kinase A inhibitor alisertib (0.2 µM), or the topoisomerase II inhibitor etoposide (0.5 µM). Cells were treated for with inducers for 4 days (refreshed on day 2) with drug withdrawn on day 4 to allow the drug to be exhausted and senescent-like phenotype is established after day 6. Induction was confirmed by senescence-associated β-galactosidase (SA-β-gal) activity, p16 or p21 immunostaining, and absence of proliferation (Ki67-negative).

### Senescence-Associated Beta-Galactosidase Staining

SA-β-gal activity was detected at pH 6.0 by a standard chromogenic assay and quantified as the percentage of β-gal-positive cells across multiple fields per condition using Fiji software. Senescence β-Galactosidase Staining Kit (CST #9860) was used according to standard protocol.

### Multiplex and Sequential Immunofluorescence

Sequential multiplex immunofluorescence staining was performed on 5-µm FFPE tissue sections following antigen retrieval using a commercial antigen unmasking solution (Vector Laboratories, H-3301-250). Standard immunofluorescence staining was carried out using a blocking and antibody dilution buffer consisting of PBS supplemented with 3% BSA and 0.3% Triton X-100. After image acquisition, antibody stripping was performed by incubating slides in stripping buffer (62.5 mM Tris-HCl, pH 6.8, 2% w/v SDS, and 114.4 mM β-mercaptoethanol in distilled water) at 56°C for 30 minutes. Subsequent rounds of staining, imaging, and stripping were repeated sequentially for multiplex marker detection. Images from all staining cycles were aligned, merged, and analyzed using Fiji software. The following antibodies were used, KRT19 (A53-B/A2, Thermo MAB-12663, for human; DSHB #TROMA-III for mouse), αSMA (Abcam #ab124964; ab7817-500), CD45 (Abcam #ab10558), p16 (Roche #805-4713 or Leica #PA0016 for human; Abcam # ab232402/211542 for mouse), p21 (CST #2947 for Human; Abcam #188224 for mouse), Ki67 (BD #561126 for human; Thermo #14-5698-82 for mouse), γH2A.X (CST #9718). The Alexa fluor labelled Donkey anti-mouse/rabbit/rat/goat secondary antibodies were purchased from ThermoFisher. Nuclei were counterstained with Hoechst 33342.

### Telomere-Associated Foci (TAF)

Telomere-associated foci were detected by combining γH2A.X immunofluorescence with telomere fluorescence in situ hybridization and scored as colocalized foci per nucleus.^33^ Briefly, FFPE slides were antigen retrieved and immunostained with γH2A.X as described above, then fixed with 4% PFA for 10 min at RT and dehydrated by series graded icy-ethanol. Tissue sections were denatured at 82°C for 10 min in hybridization buffer consisting of 70% formamide, 25 mM MgCl₂, 0.1 M Tris-HCl (pH 7.2), and 5% Roche blocking reagent, containing 2.5 μg/mL Cy3-labeled telomere-specific PNA probe (Panagene). Hybridization was carried out for 2 h at room temperature in a humidified dark chamber. After hybridization, sections were washed once in 70% formamide/2× SSC for 10 min, followed by two 10-min washes in 2× SSC and one 10-min wash in PBS. Sections were imaged using in-depth Z stacking (a minimum of 40 optical slices with 63× objective) followed by Huygens (SVI) deconvolution.

### Single-Nucleus RNA Sequencing

Nuclei were isolated from human CCA tumors (n=10 patients) and processed for single-nucleus RNA sequencing on the 10x Genomics Chromium platform. Reads were aligned to the human reference, and nuclei were quality-filtered, normalized, integrated, and clustered. Cancer-cell clusters were annotated using KRT19 and epithelial lineage markers, and a senescence-high cancer-cell cluster was identified by senescence-program enrichment.

### Senescence Transcriptional Scoring

Senescence and senescence-associated secretory phenotype programs (SenMayo,^34^ Fridman senescence,^35^ and Reactome cellular senescence^36^) were scored per cell using UCell. Single-cell expression comparisons used mixed-effects models with patient as a random effect to avoid pseudoreplication.

### Bulk RNA Sequencing

Bulk RNA sequencing was performed on the Mayo CCA cohort and an external Asian cohort (OEP00105), and senescence transcriptional scores were computed per sample by single-sample gene-set scoring. For macrophage profiling, RNA was extracted from BMDMs after conditioned-medium treatment, sequenced, and analyzed for differential gene expression relative to control.

### Spatial Transcriptomics

Xenium spatial transcriptomics was performed on FFPE human CCA sections using a probe set that included *CDKN2A* and *GDF15*. *GDF15*-expressing cancer cells and neighboring macrophages were identified, and TAM immunosuppression scores were computed as a function of proximity to *GDF15*-expressing cancer cells. GeoMx digital spatial profiling was used to relate *GDF15*-positive cancer-cell density to TAM proximity.

### Quantitative RT-PCR

Total RNA was extracted using Trizol and reverse-transcribed using LunaScript® RT SuperMix (NEB # M3010L), and target transcripts were quantified by quantitative PCR using Luna Universal qPCR Master Mix (NEB # M3003X) on LightCycler® PRO system using the following primers, *tGFP* (F: AGGACAGCGTGATCTTCACC, R: CTTGAAGTGCATGTGGCTGT), *hGDF-15* (F: GACCCTCAGAGTTGCACTCC, R: GCCTGGTTAGCAGGTCCTC), *Gdf-15* (F: AGCCGAGAGGACTCGAACTCAG, R, GGTTGACGCGGAGTAGCAGCT), *Arginase 1* (*Arg 1*, F: CTCCAAGCCAAAGTCCTTAGAG, R: GGAGCTGTCATTAGGGACATCA), *Nos2* (*iNos*, F: GTTCTCAGCCCAACAATACAAGA, R: GTGGACGGGTCGATGTCAC), *Opn* (*Spp1*, F: CAGCCTGCACCCAGATCCTA, R: GCGCAAGGAGATTCTGCTTCT), *Cxcl9* (F: GGAGTTCGAGGAACCCTAGTG, R: GGGATTTGTAGTGGATCGTGC), *Tgfbr2* (F, CCTACTCTGTCTGTGGATGACC, R: GACATCCGTCTGCTTGAACGAC), *hGAPDH* (F: GGAGCGAGATCCCTCCAAAAT, R: GGCTGTTGTCATACTTCTCATGG), *Gapdh* (F: AGGTCGGTGTGAACGGATTTG, R:TGTAGACCATGTAGTTGAGGTCA) or *Actb* (F: AATCGTGCGTGACATCAAAGAG, R: GCCATCTCCTGCTCGAAGTC) were used as housekeeping genes for calculating relative gene expression by the comparative Ct method.

### Survival Analysis

Overall survival was modeled by multivariable Cox regression adjusting for age, sex, and stage, with the same model specification applied independently in the Mayo discovery cohort (n=158, 108 events) and the external Asian cohort (OEP00105, n=244, 99 events) (patient characteristics in Supplementary Table S1); multivariable Cox models included only cases with complete age, sex, and stage data. Kaplan-Meier survival with the log-rank test was used for score-high versus score-low comparisons in the Mayo cohort (n=233).

### Cytokine Array

Conditioned medium from proliferating or Sen-L human RBE cells (doxorubicin- or gemcitabine/cisplatin-induced) was profiled with the Proteome Profiler Human XL Cytokine Array (R&D Systems #ARY022B) according to the standard manufacturer’s protocol except being developed with SuperSignal West Femto Maximum Sensitivity Substrate (ThermoFisher #34096) and visualized using BIO-RAD ChemiDoc MP imager. Signals were quantified by densitometry using Fiji software and ranked to identify the dominant secreted factor.

### Immunoblotting

Macrophage lysates in RIPA buffer supplemented with proteinase inhibitor and phosphatase inhibitor cocktails were resolved by SDS-PAGE, transferred to nitrocellulose membranes, and probed for phospho-Stat6 (CST #56554S), Stat6 (CST #5397S), phospho-Stat3 (CST #9145S), Stat3 (CST #9139S), phospho-SMAD3 (CST #9520T) and SMAD3 (CST #9523T) at 1: 1000 in TBST with 5% BSA. Immun-Star Goat Anti-Mouse/Rabbit (GAM)-HRP Conjugate (BIO-RAD #1705047/1705046) were used. Signal was developed with SuperSignal West Femto Maximum Sensitivity Substrate (ThermoFisher #34096) and visualized using BIO-RAD ChemiDoc MP imager.

### GDF-15 Proximity Labeling and TGFBR2 Identification

Candidate macrophage GDF-15 receptors were identified by proximity-dependent biotinylation using a secreted GDF-15-BioID2 fusion, with a SEC61B-BioID2 fusion as a spatial control, followed by streptavidin enrichment and anti-TGFBR2 immunoblot. TGFBR2 dependence was tested by *Tgfbr2* loss-of-function in macrophages.

### Recombinant GDF-15

Carrier-free recombinant GDF-15 was used for macrophage stimulation, with each lot confirmed bioactive and free of TGF-β1. Human recombinant GDF-15 was from Qkine (Qk017) and murine recombinant GDF-15 from R&D Systems (8944-GD/CF).

### Flow Cytometry

Under deep anesthesia using 1.5–3% isoflurane, the spleen, tumor, and tumor-adjacent liver were excised and weighed. Tumors were finely chopped with scissors in C tubes, dissociated into single-cell suspensions with tissue-specific dissociation kits (Miltenyi #130-096-730) on a gentleMACS Octo Dissociator (Miltenyi Biotec) and filtered through a 70-µm strainer. CD45 positive cells were enriched with CD45 (TIL) MicroBeads, mouse (Miltenyi # 130-110-618) according to the manufacturer’s instruction. Single-cell suspensions were stained with a Fixable Viability Stain 510 (BD #564406, 1: 1000) or 660 (BD #564405, 1: 2000) for 20 min at room temperature (RT) in PBS, and surface-labeled with indicated antibodies in FACS buffer (PBS with 10% FBS and 1 mM EDTA) for 30 min at 4 C, then fixed (1h at RT) and stained with intracellular markers using True-Nuclear Transcription Factor Buffer Set (Biolegend #424401) for 1h at RT. The following antibodies were used (1:100 dilution) for flow cytometry analysis: CD11b_PerCP-Cy5.5 (Biolegend #101228), CD11c_BV650 (ThermoFisher #17-0114-82), F4/80_BV421 (Biolegend #123132), Ly6C_APC-Cy7 (Biolegend #128026), Ly6G_AF488 (Biolegend #127626), CD206_PE-Cy7 (Biolegend #141720), CD3_APC-Cy7 (Biolegend #100222), CD4_PerCP-Cy5.5 (Biolegend #116012), CD8a_BV421 (BD #563898), NK1.1_PE (ThermoFisher #12-5941-83), CD19_BV711 (ThermoFisher #407-0193-82), FoxP3_APC (ThermoFisher #17-5773-82), Granzyme B_PE-Cy7 (Biolegend #372214). Data were acquired on a MACSQuant Analyzer (Miltenyi) or ZE5 Cell Analyzer (BIO-RAD) and analyzed in FlowJo software.

### Statistics

The majority of experiments were repeated at least twice to obtain robust data for the indicated statistical analyses. For all readouts, examiners were blinded. The experiments (mice per group) were randomized. Data are presented as mean +/- SEM. Pairwise comparisons used the two-tailed t test or Mann-Whitney test; multigroup comparisons used one-way ANOVA or Kruskal-Wallis with correction for multiple comparisons; survival was analyzed by Kaplan-Meier with the log-rank test and by multivariable Cox regression. Single-cell expression comparisons used mixed-effects models with patient as a random effect. Animals were randomized to treatment, with group allocation matched for tumor size where applicable. GraphPad Prism 11 was used for statistical and visual output. Human data analyses were performed using R (v4.2.1) on RStudio (v2022.07.0). P-value < 0.05 indicates statistical significance.

## ACKNOWLEDGEMENTS

We thank the Mayo Clinic shared resources and cores that supported this work, including the NIDDK-funded Mayo Clinic Digestive Disease Research Core Center (DDRCC; P30DK84567), the Mayo Clinic Hepatobiliary SPORE, the Mayo Clinic Comprehensive Cancer Center and its shared resources, and the Mayo Clinic Hepatobiliary Biorepository.

## Abbreviations

ANOVA: analysis of variance
BMDM: bone marrow-derived macrophage
CCA: cholangiocarcinoma
CFSE: carboxyfluorescein succinimidyl ester
CTL: cytotoxic T lymphocyte
DAPI: 4′,6-diamidino-2-phenylindole
DMEM: Dulbecco’s Modified Eagle Medium
FACS: fluorescence-activated cell sorting
FFPE: formalin-fixed paraffin-embedded
GDF-15: growth differentiation factor 15
Gem/Cis: gemcitabine/cisplatin
IACUC: Institutional Animal Care and Use Committee
IRB: Institutional Review Board
M-CSF: macrophage colony-stimulating factor
MDSC: myeloid-derived suppressor cell
M-MDSC: monocytic myeloid-derived suppressor cell
NK: natural killer
PD-1: programmed cell death protein 1
PD-L1: programmed death-ligand 1
PMN-MDSC: polymorphonuclear myeloid-derived suppressor cell
qPCR: quantitative PCR
SA-β-gal: senescence-associated β-galactosidase
SASP: senescence-associated secretory phenotype
SDS-PAGE: sodium dodecyl sulfate-polyacrylamide gel electrophoresis
SEM: standard error of the mean
Sen-L: senescent-like
shRNA: short hairpin RNA
siRNA: small interfering RNA
TAF: telomere-associated foci
TAM: tumor-associated macrophage
TGF-β: transforming growth factor-β
tGFP: turbo green fluorescent protein
TiME: tumor immune microenvironment
Treg: regulatory T cell
UMAP: uniform manifold approximation and projection

---

**Supplementary Figure S1.**
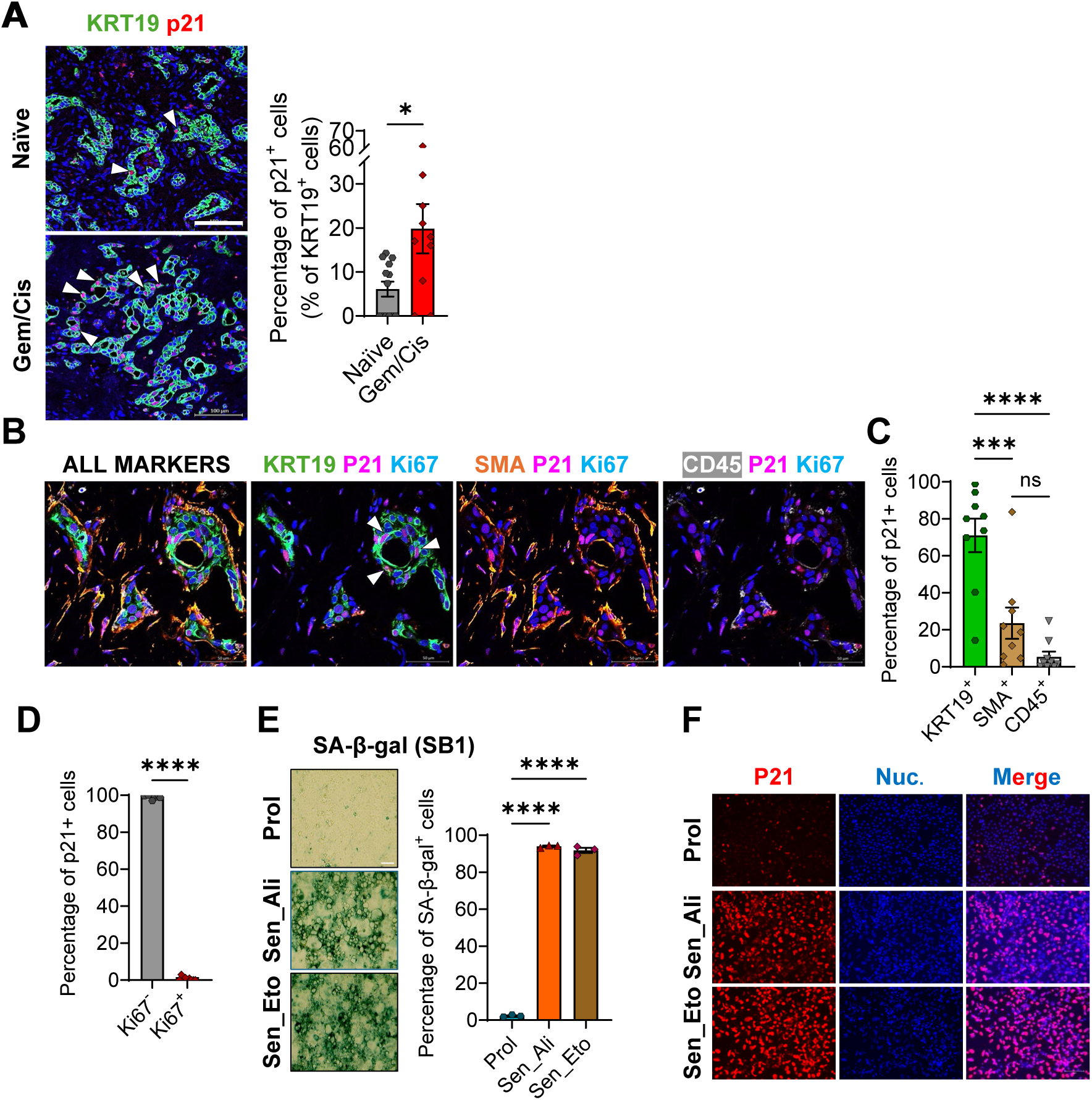
(A) p21^+^ cancer cells in treatment-naive (n=15) versus Gem/Cis-treated (n=19) human CCA. (B-D) Sequential immunofluorescence of p21^+^ cancer cells (KRT19^+^), cancer-associated fibroblasts (α-SMA^+^), and immune cells (CD45^+^), and Ki67 status. (E-F) Senescence induction in SB1 cells with alisertib (0.2 µM, Aurora kinase A inhibitor) or etoposide (0.5 µM, topoisomerase II inhibitor) over 6 days, with drug withdrawn at day 4, confirmed by SA-β-gal and (E) p21 immunostaining (F).

**Supplementary Figure S2.**
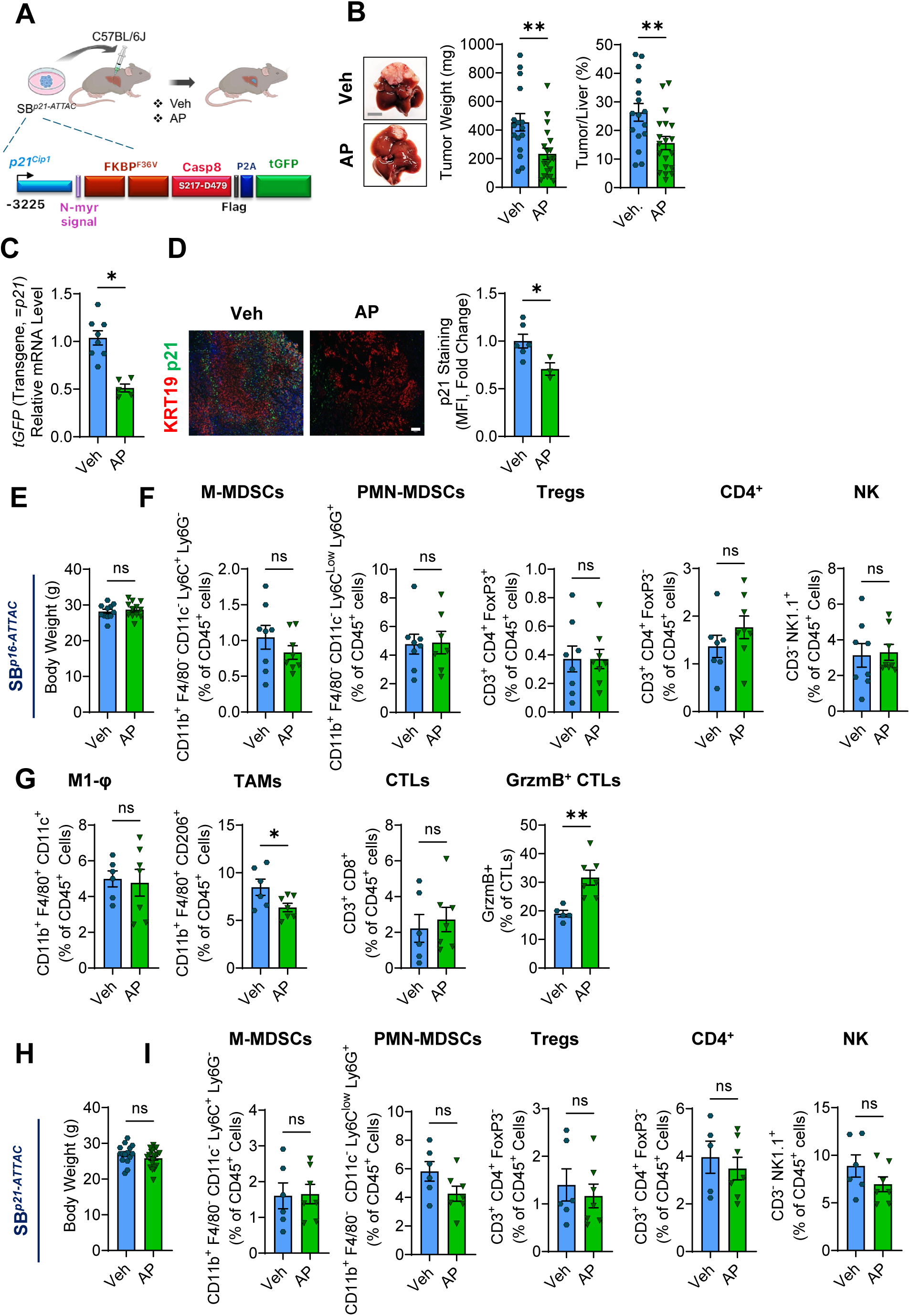
(A) SB1 *p21-ATTAC* in vivo study schematic. (B) Representative gross images and tumor-burden quantification, vehicle versus AP-treated SB1 *p21-ATTAC* tumors. (C) Relative mRNA of p21-promoter tGFP reporter in veh and AP treated murine SB1 *p21-ATTAC* tumors. (D) p21 mean fluorescence intensity in veh and AP treated murine SB1 *p21-ATTAC* tumors. Scale bar, 100 µm. (E) Body weights of SB1 *p16-ATTAC* tumor bearing mice treated with or AP. (F) Monocytic myeloid-derived suppressor cell (M-MDSC), polymorphonuclear myeloid-derived suppressor cell (PMN-MDSC), regulatory T-cell (Treg), CD4^+^ T cells, and natural killer (NK) frequencies (% of CD45^+^ cells) in in SB1 *p16-ATTAC* tumors. (G) M1 macrophage, TAM, CD8^+^ CTL, granzyme B^+^ CD8^+^ CTL frequencies in SB1 *p21-ATTAC* tumors. (H) Body weights of SB1 *p21-ATTAC* tumor bearing mice treated with or AP. (I) M-MDSC, PMN-MDSC, Treg, CD4^+^ T-cell, and NK-cell frequencies in SB1 *p21-ATTAC* tumors. Data are mean ± SEM; *, P<0.05; **, P<0.01.

**Supplementary Figure S3.**
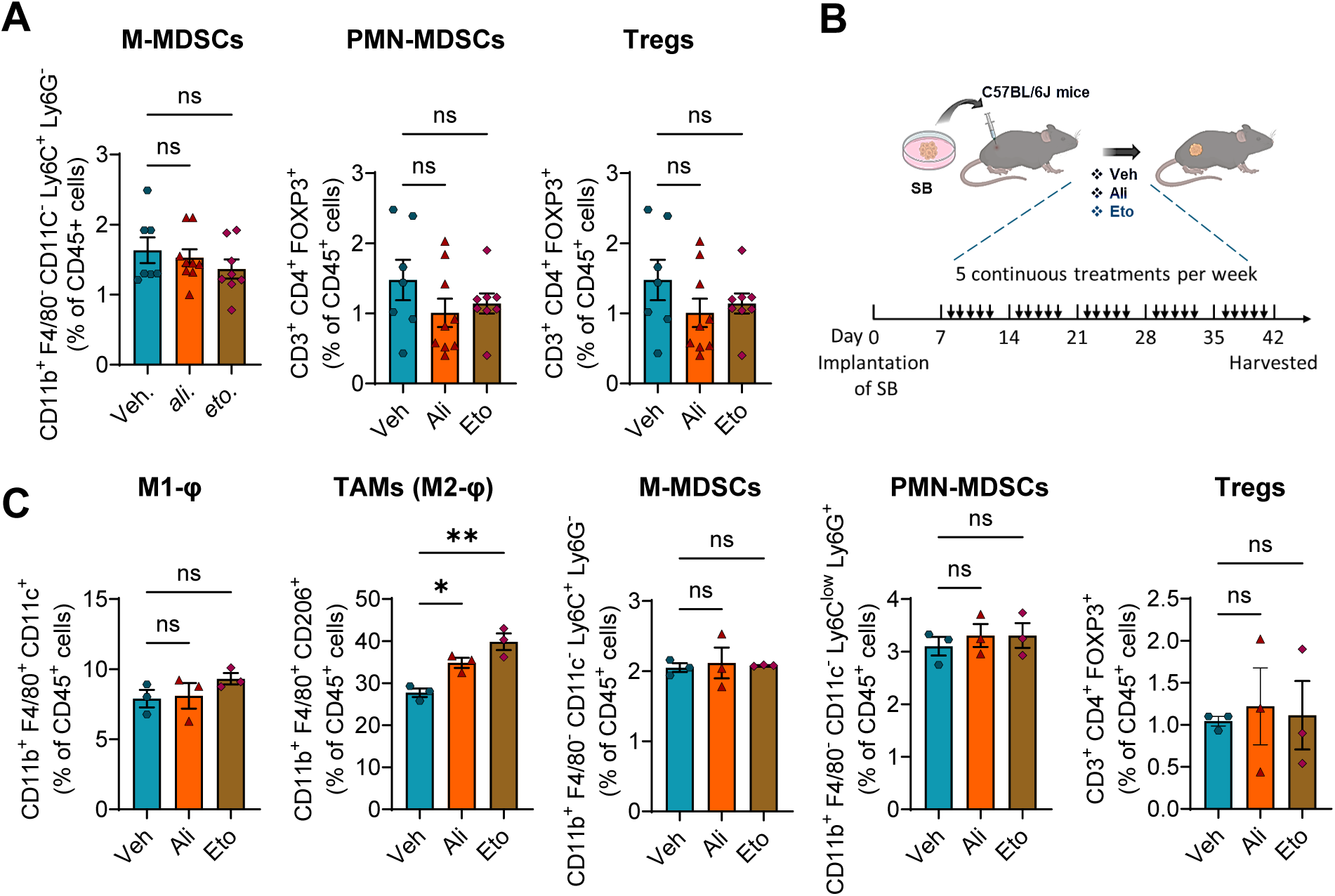
(A) M-MDSC, PMN-MDSC, and regulatory T-cell (Treg) frequencies (% of CD45^+^ cells) in the SB1 tumor bearing mice treated with vehicle, alisertib, or etoposide. (B) Schematic of subcutaneous tumor model study. (C) M1 macrophage, TAM, M-MDSC, PMN-MDSC, Treg frequencies in subcutaneous SB1 tumor bearing mice treated with vehicle, alisertib, or etoposide. Data are mean ± SEM; *, P<0.05; **; ns, not significant.

**Supplementary Figure S4.**
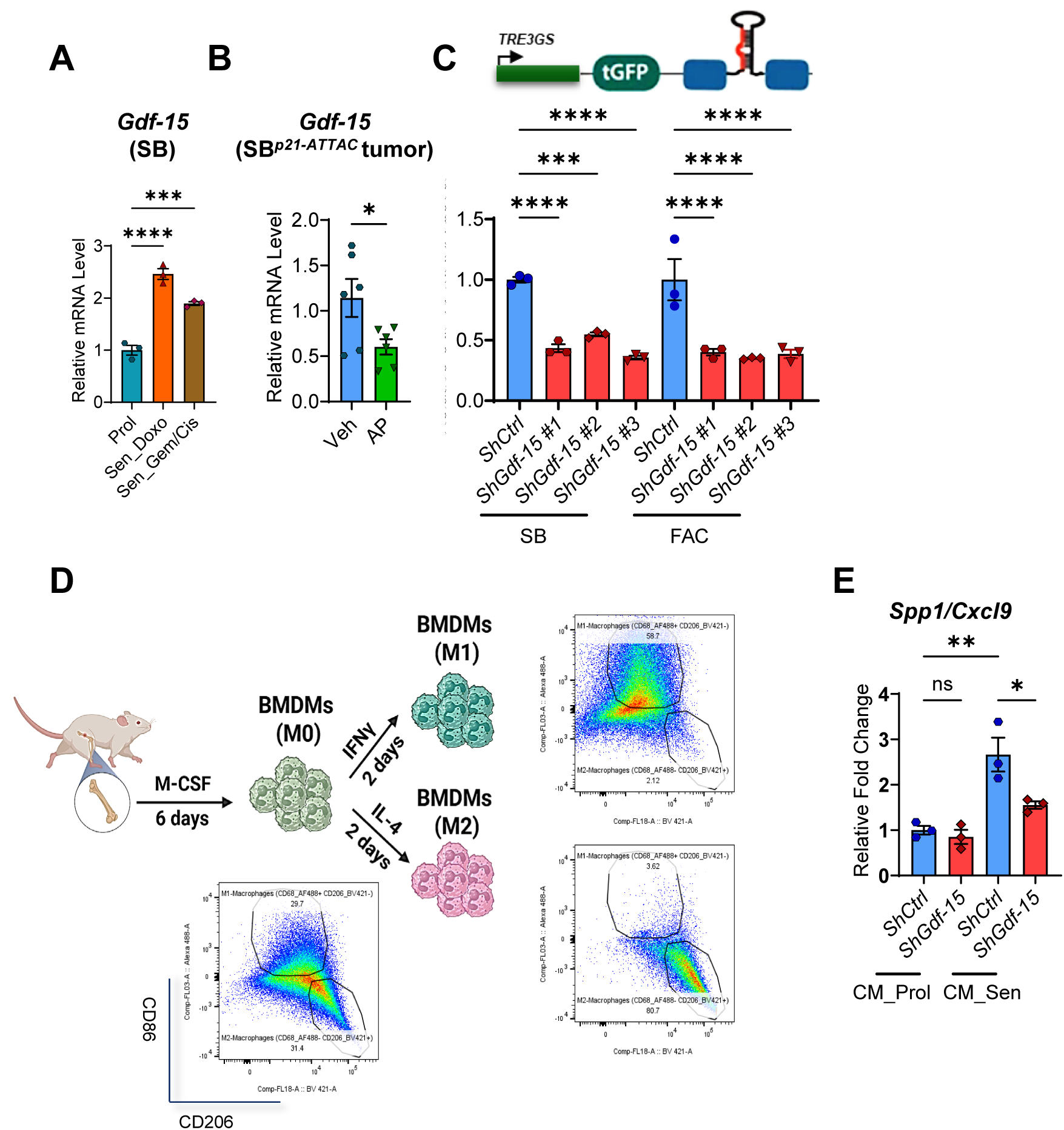
(A) *Gdf15* mRNA in SB1 cells treated with vehicle (proliferating), doxorubicin (Dox) or gemcitabine/cisplatin (Gem/Cis). (B) Intratumoral *Gdf15* mRNA in *p21-ATTAC* murine tumors treated with vehicle or AP. (C) Schematic and qPCR validation of *Gdf15*-targeting shRNAs. (D) Schematic of in vitro studies. (E) *Arg1* and *iNOS* mRNA in bone-marrow-derived macrophage (BMDM) incubated for 72 h with conditioned medium from vehicle- or alisertib-treated SB1 cells expressing *shCtrl* or *shGdf15*. Data are mean ± SEM; *, P<0.05; ***, P<0.001; ****, P<0.0001.

**Supplementary Figure S5.**
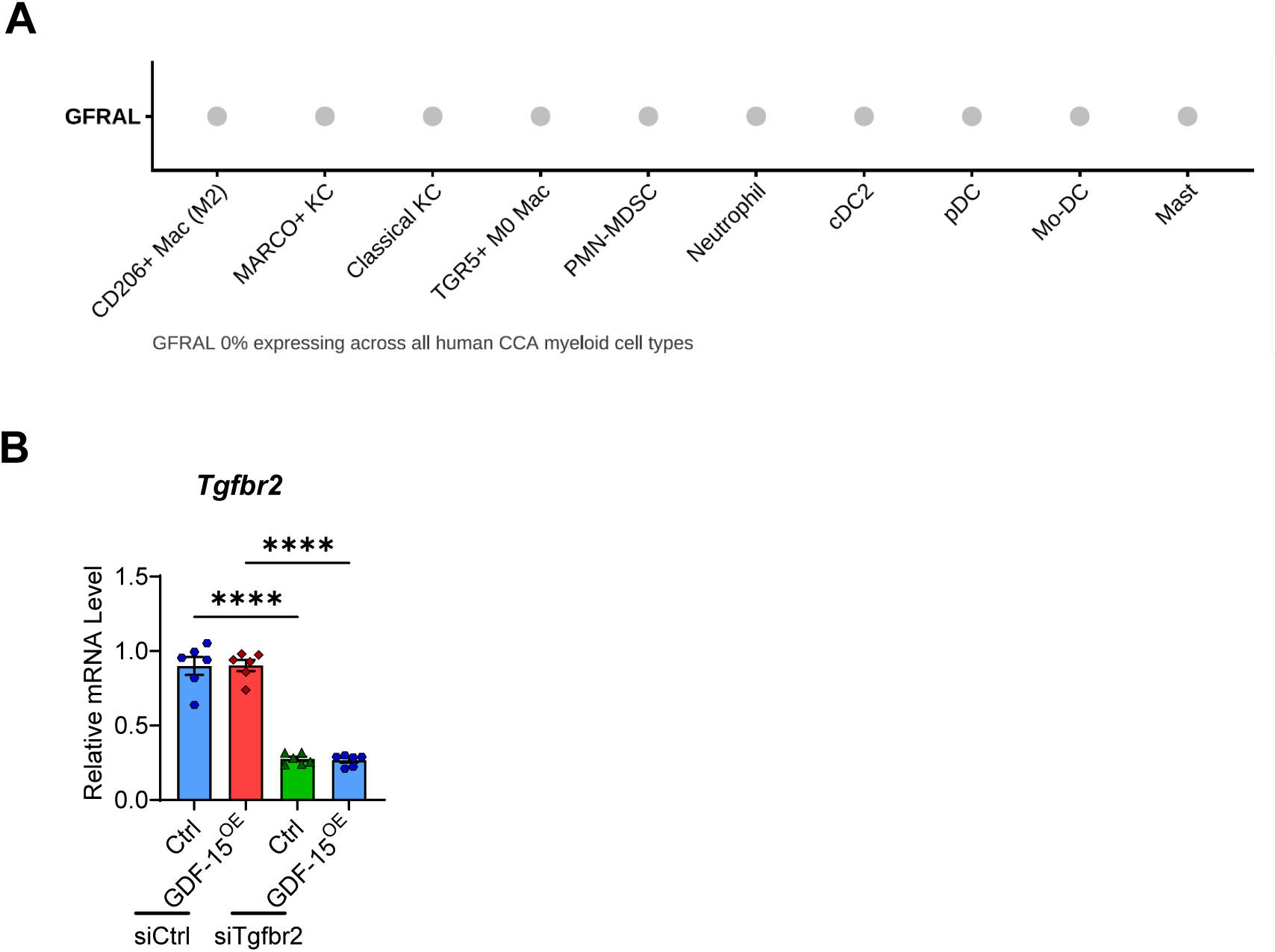
(A) Single-cell RNA-seq dot plot of *GFRAL* across CD206⁺ M2-like macrophages, MARCO⁺ and classical Kupffer cells (KC), TGR5⁺ M0 macrophages, polymorphonuclear myeloid-derived suppressor cells (PMN-MDSCs), neutrophils, conventional type 2 dendritic cells (cDC2), plasmacytoid dendritic cells (pDC), monocyte-derived dendritic cells (Mo-DC), and mast cells. *GFRAL* was undetectable (0% of cells expressing) in all populations. Dot size and color denote the fraction of cells expressing and mean scaled expression, respectively. (B) *Tgfbr2* mRNA in BMDMs transfected with *siCtrl* or *siTgfbr2* and stimulated with control or rGDF-15. n = 6. Data are mean ± SEM; *, P<0.05; **, P<0.01; ***, P<0.001; ****, P<0.0001; ns, not significant.

**Supplementary Figure S6.**
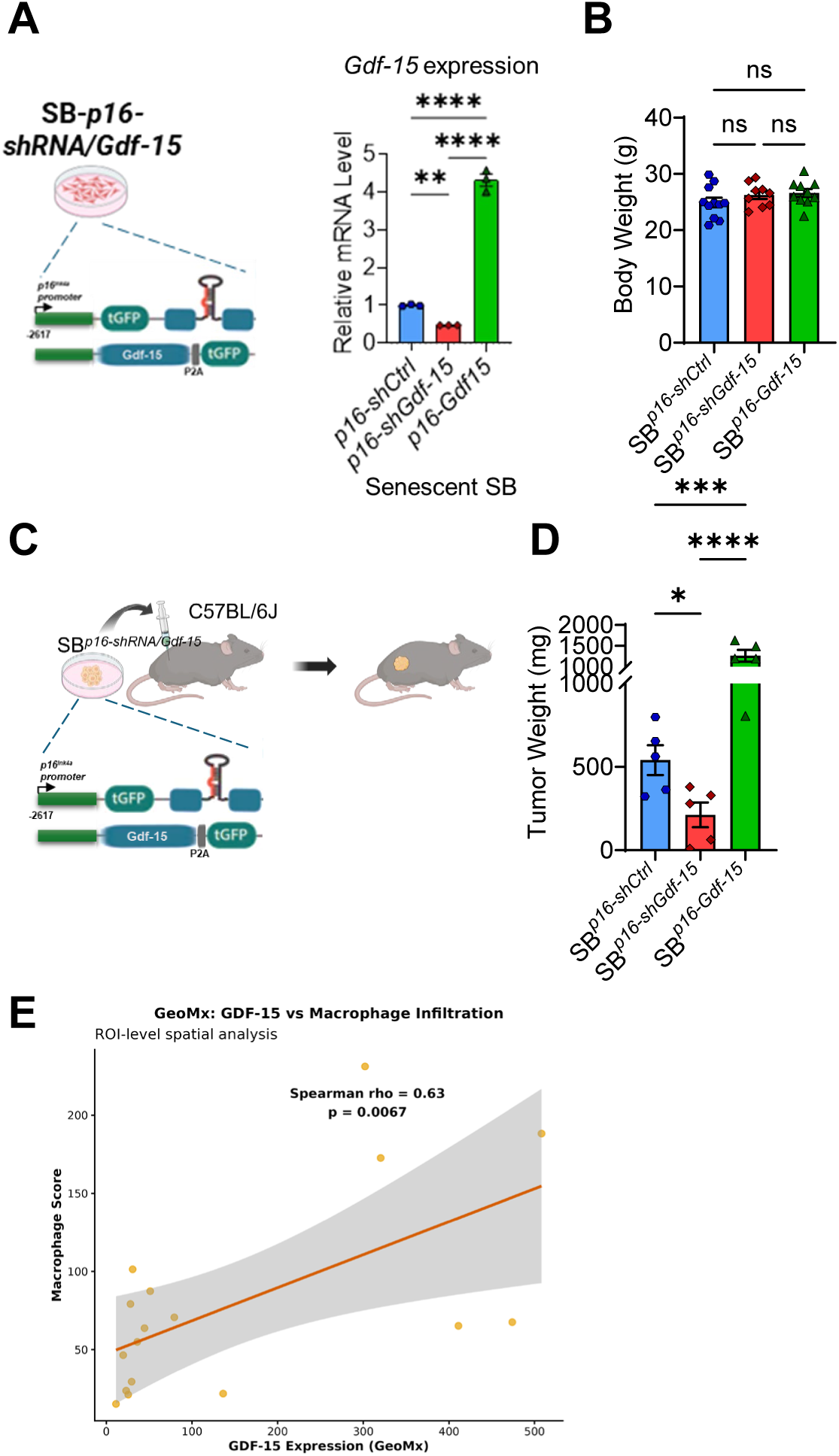
(A) Strategy for p16-promoter-restricted *Gdf15* silencing (shRNA) and *Gdf15* overexpression. (B) qPCR validation of *Gdf15* knockdown and overexpression in senescent-like SB1 cells. (C-D) Subcutaneous model and tumor-burden quantification. (E) GeoMx spatial profiling of GDF-15 expression and macrophage infiltration in human CCA. Region-of-interest (ROI)-level analysis plotting GDF-15 expression against macrophage score; each point is one ROI, with the linear fit and 95% confidence band shown. Spearman ρ = 0.63, *P* = 0.0067. Data are mean ± SEM; **, P<0.01; ****, P<0.0001.

**Supplementary Figure S7.**
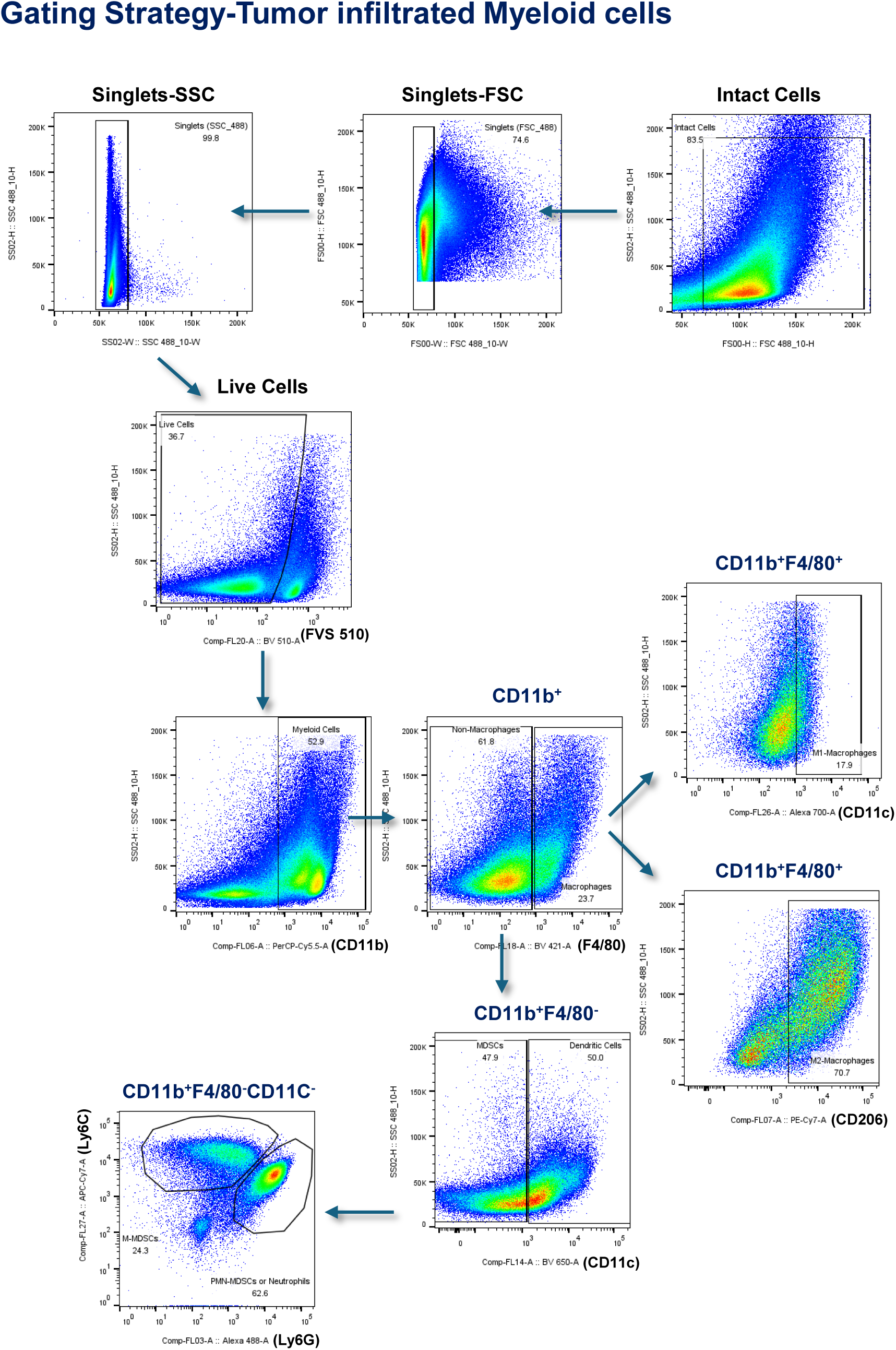
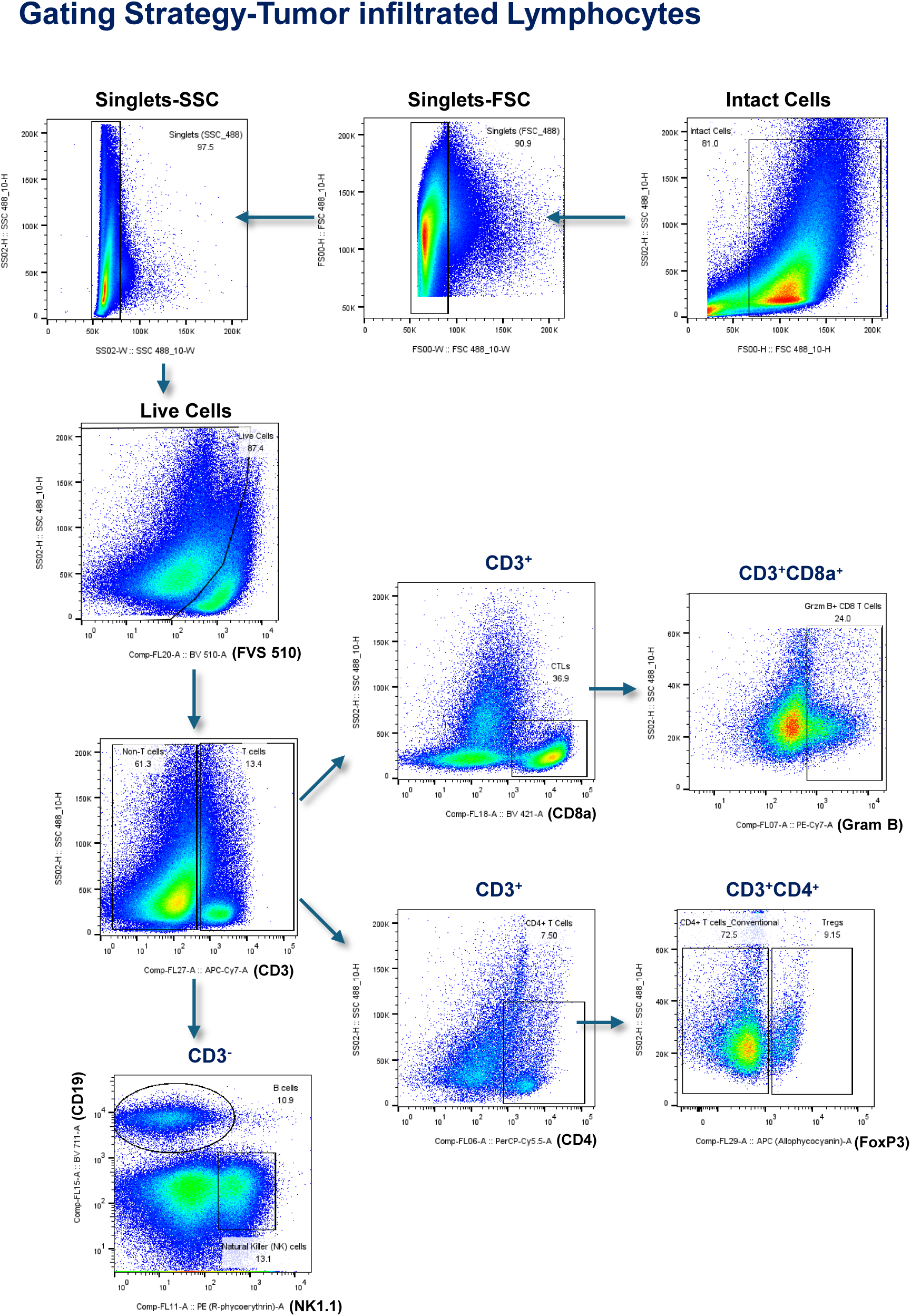
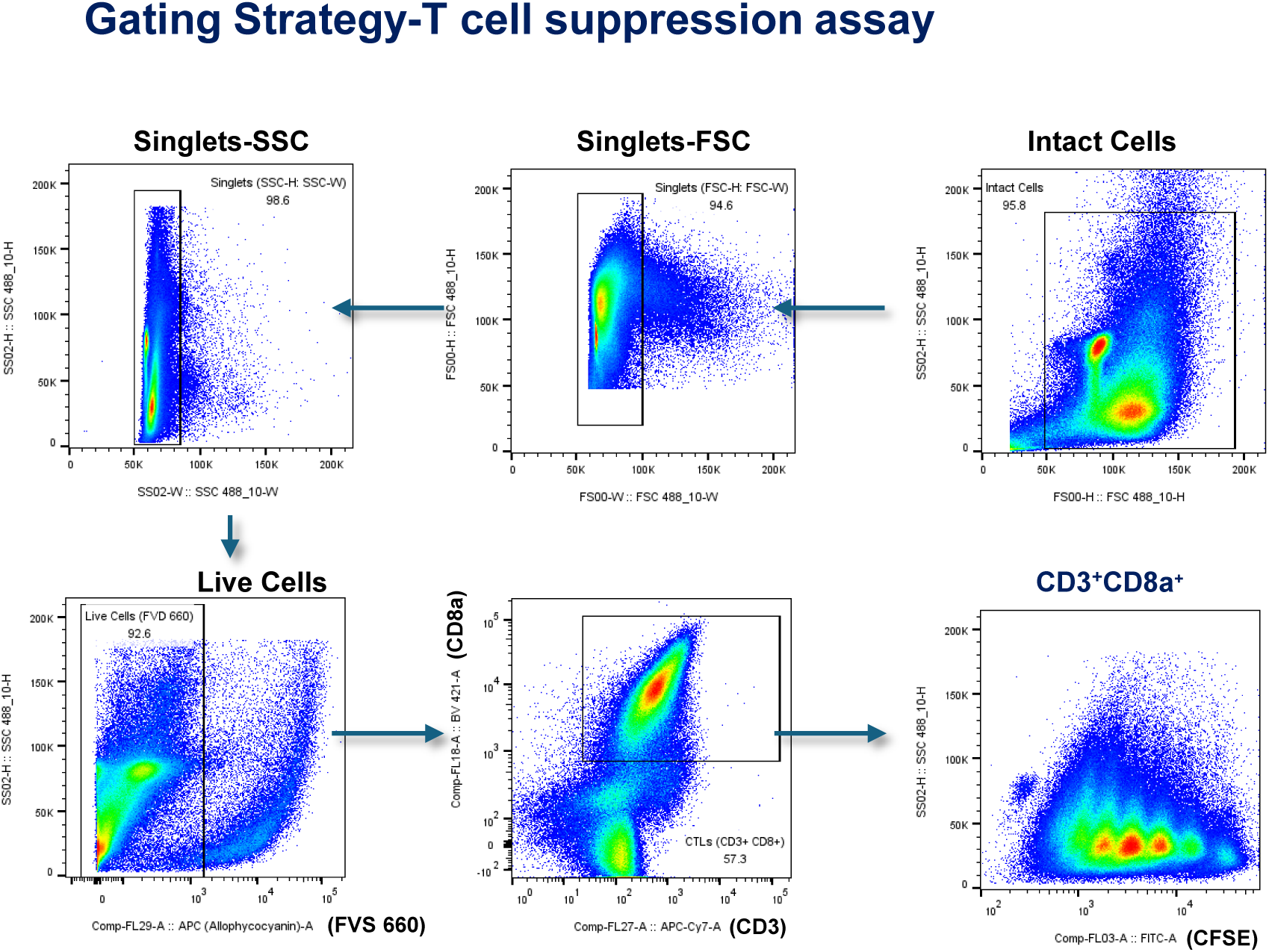
Gating strategies for flow cytometry used in immunoprofiling of tumor-infiltrated CD45^+^ cells, and T cell suppression assay.

## REFERENCES

1. Ilyas SI, Affo S, Goyal L, Lamarca A, Sapisochin G, Yang JD, et al. Cholangiocarcinoma -novel biological insights and therapeutic strategies. Nat Rev Clin Oncol 2023;20:470–86.

2. Oh DY, Ruth He A, Qin S, Chen LT, Okusaka T, Vogel A, et al. Durvalumab plus gemcitabine and cisplatin in advanced biliary tract cancer. NEJM Evid 2022;1:EVIDoa2200015.

3. Kelley RK, Ueno M, Yoo C, Finn RS, Furuse J, Ren Z, et al. Pembrolizumab in combination with gemcitabine and cisplatin compared with gemcitabine and cisplatin alone for patients with advanced biliary tract cancer (KEYNOTE-966): A randomised, double-blind, placebo-controlled, phase 3 trial. Lancet 2023;401:1853–65.

4. Vasan N, Baselga J, Hyman DM. A view on drug resistance in cancer. Nature 2019;575:299–309.

5. Demaria M, O’Leary MN, Chang J, Shao L, Liu S, Alimirah F, et al. Cellular senescence promotes adverse effects of chemotherapy and cancer relapse. Cancer Discov 2017;7:165–76.

6. Duy C, Li M, Teater M, Meydan C, Garrett-Bakelman FE, Lee TC, et al. Chemotherapy induces Senescence-Like resilient cells capable of initiating AML recurrence. Cancer Discov 2021;11:1542–61.

7. Chaib S, López-Domínguez JA, Lalinde-Gutiérrez M, Prats N, Marin I, Boix O, et al. The efficacy of chemotherapy is limited by intratumoral senescent cells expressing PD-L2. Nat Cancer 2024;5:448–62.

8. Maggiorani D, Le O, Lisi V, Landais S, Moquin-Beaudry G, Lavallée VP, et al. Senescence drives immunotherapy resistance by inducing an immunosuppressive tumor microenvironment. Nat Commun 2024;15:2435.

9. Saleh T, Tyutyunyk-Massey L, Gewirtz DA. Tumor cell escape from Therapy-Induced senescence as a model of disease recurrence after dormancy. Cancer Res 2019;79:1044–6.

10. Marin I, Boix O, Garcia-Garijo A, Sirois I, Caballe A, Zarzuela E, et al. Cellular senescence is immunogenic and promotes antitumor immunity. Cancer Discov 2023;13:410–31.

11. Chen H, Ho Y, Mezzadra R, Adrover JM, Smolkin R, Zhu C, et al. Senescence rewires microenvironment sensing to facilitate antitumor immunity. Cancer Discov 2023;13:432–53.

12. Coppé JP, Desprez PY, Krtolica A, Campisi J. The senescence-associated secretory phenotype: The dark side of tumor suppression. Annu Rev Pathol 2010;5:99–118.

13. Schmitt CA, Wang B, Demaria M. Senescence and cancer -role and therapeutic opportunities. Nat Rev Clin Oncol 2022;19:619–36.

14. Tomlinson JL, Valle JW, Ilyas SI. Immunobiology of cholangiocarcinoma. J Hepatol 2023;79:867–75.

15. Loeuillard E, Yang J, Buckarma E, Wang J, Liu Y, Conboy C, et al. Targeting tumor-associated macrophages and granulocytic myeloid-derived suppressor cells augments PD-1 blockade in cholangiocarcinoma. J Clin Invest 2020;130:5380–96.

16. Greten TF, Schwabe R, Bardeesy N, Ma L, Goyal L, Kelley RK, et al. Immunology and immunotherapy of cholangiocarcinoma. Nat Rev Gastroenterol Hepatol 2023;20:349–65.

17. Ruffolo LI, Jackson KM, Kuhlers PC, Dale BS, Figueroa Guilliani NM, Ullman NA, et al. GM-CSF drives myelopoiesis, recruitment and polarisation of tumour-associated macrophages in cholangiocarcinoma and systemic blockade facilitates antitumour immunity. Gut 2022;71:1386–98.

18. Zhou L, DeMarco KD, Murphy KC, Wu Z, Li J, Johnson C, et al. p21-Positive senescent stromal cells promote prostate cancer immune suppression and progression that can be reversed by senolytic therapy. Cancer Discov 2026;16:571–91.

19. Belle JI, Sen D, Baer JM, Liu X, Lander VE, Ye J, et al. Senescence defines a distinct subset of myofibroblasts that orchestrates immunosuppression in pancreatic cancer. Cancer Discov 2024;14:1324–55.

20. Ye J, Baer JM, Faget DV, Morikis VA, Ren Q, Melam A, et al. Senescent CAFs mediate immunosuppression and drive breast cancer progression. Cancer Discov 2024;14:1302–23.

21. Yasuda T, Koiwa M, Yonemura A, Miyake K, Kariya R, Kubota S, et al. Inflammation-driven senescence-associated secretory phenotype in cancer-associated fibroblasts enhances peritoneal dissemination. Cell Rep 2021;34:108779.

22. Prieto LI, Sturmlechner I, Graves SI, Zhang C, Goplen NP, Yi ES, et al. Senescent alveolar macrophages promote early-stage lung tumorigenesis. Cancer Cell 2023;41:1261–1275 e6.

23. Wischhusen J, Melero I, Fridman WH. Growth/Differentiation Factor-15 (GDF-15): From biomarker to novel targetable immune checkpoint. Front Immunol 2020;11:951.

24. Melero I, de Miguel Luken M, de Velasco G, Garralda E, Martín-Liberal J, Joerger M, et al. Neutralizing GDF-15 can overcome anti-PD-1 and anti-PD-L1 resistance in solid tumours. Nature 2025;637:1218–27.

25. Evans DS, Young D, Tanaka T, Basisty N, Bandinelli S, Ferrucci L, et al. Proteomic analysis of the Senescence-Associated secretory phenotype: GDF-15, IGFBP-2, and Cystatin-C are associated with multiple aging traits. J Gerontol A Biol Sci Med Sci 2024;79(3).

26. Basisty N, Kale A, Jeon OH, Kuehnemann C, Payne T, Rao C, et al. A proteomic atlas of senescence-associated secretomes for aging biomarker development. PLoS Biol 2020;18:e3000599.

27. Haake M, Haack B, Schäfer T, Harter PN, Mattavelli G, Eiring P, et al. Tumor-derived GDF-15 blocks LFA-1 dependent T cell recruitment and suppresses responses to anti-PD-1 treatment. Nat Commun 2023;14:4253.

28. Hsu J, Crawley S, Chen M, Ayupova DA, Lindhout DA, Higbee J, et al. Non-homeostatic body weight regulation through a brainstem-restricted receptor for GDF15. Nature 2017;550:255–9.

29. Hes C, Gui LT, Bay A, Alvarez F, Katz P, Paul T, et al. GDNF family receptor alpha-like (GFRAL) expression is restricted to the caudal brainstem. Mol Metab 2025;91:102070.

30. de Jager SCA, Bermúdez B, Bot I, Koenen RR, Bot M, Kavelaars A, et al. Growth differentiation factor 15 deficiency protects against atherosclerosis by attenuating CCR2-mediated macrophage chemotaxis. J Exp Med 2011;208:217–25.

31. Artz A, Butz S, Vestweber D. GDF-15 inhibits integrin activation and mouse neutrophil recruitment through the ALK-5/TGF-βRII heterodimer. Blood 2016;128:529–41.

32. Gil J. The challenge of identifying senescent cells. Nat Cell Biol 2023;25:1554–6.

33. Hewitt G, Jurk D, Marques FDM, Correia-Melo C, Hardy T, Gackowska A, et al. Telomeres are favoured targets of a persistent DNA damage response in ageing and stress-induced senescence. Nat Commun 2012;3:708.

34. Saul D, Kosinsky RL, Atkinson EJ, Doolittle ML, Zhang X, LeBrasseur NK, et al. A new gene set identifies senescent cells and predicts senescence-associated pathways across tissues. Nat Commun 2022;13:4827.

35. Fridman AL, Tainsky MA. Critical pathways in cellular senescence and immortalization revealed by gene expression profiling. Oncogene 2008;27:5975–87.

36. Milacic M, Beavers D, Conley P, Gong C, Gillespie M, Griss J, et al. The Reactome pathway knowledgebase 2024. Nucleic Acids Res 2024;52(D1):D672–D678.

37. Tomlinson JL, Li B, Yang J, Loeuillard E, Stumpf HE, Kuipers H, et al. Syngeneic murine models with distinct immune microenvironments represent subsets of human intrahepatic cholangiocarcinoma. J Hepatol 2024;80:892–903.

38. Baker DJ, Wijshake T, Tchkonia T, LeBrasseur NK, Childs BG, van de Sluis B, et al. Clearance of p16Ink4a-positive senescent cells delays ageing-associated disorders. Nature 2011;479:232–6.

39. Baker DJ, Childs BG, Durik M, Wijers ME, Sieben CJ, Zhong J, et al. Naturally occurring p16Ink4a-positive cells shorten healthy lifespan. Nature 2016;530:184–9.

40. Yosef R, Pilpel N, Tokarsky-Amiel R, Biran A, Ovadya Y, Cohen S, et al. Directed elimination of senescent cells by inhibition of BCL-W and BCL-XL. Nat Commun 2016;7:11190.

41. Zhu Y, Tchkonia T, Fuhrmann-Stroissnigg H, Dai HM, Ling YY, Stout MB, et al. Identification of a novel senolytic agent, navitoclax, targeting the Bcl-2 family of anti-apoptotic factors. Aging Cell 2016;15:428–35.

42. Okamoto M, Koma Y, Kodama T, Nishio M, Shigeoka M, Yokozaki H. Growth differentiation factor 15 promotes progression of esophageal squamous cell carcinoma via TGF-β type II receptor activation. Pathobiology 2020;87:100–13.

43. Pang Y, Gara SK, Achyut BR, Li Z, Yan HH, Day CP, et al. TGF-β signaling in myeloid cells is required for tumor metastasis. Cancer Discov 2013;3:936–51.

44. Wang Z, He L, Li W, Xu C, Zhang J, Wang D, et al. GDF15 induces immunosuppression via CD48 on regulatory T cells in hepatocellular carcinoma. J Immunother Cancer 2021;9:e002787.

45. Ashkenazi A, Fairbrother WJ, Leverson JD, Souers AJ. From basic apoptosis discoveries to advanced selective BCL-2 family inhibitors. Nat Rev Drug Discov 2017;16:273–84.

46. Wang X, Bathina M, Lynch J, Koss B, Calabrese C, Frase S, et al. Deletion of MCL-1 causes lethal cardiac failure and mitochondrial dysfunction. Genes Dev 2013;27:1351–64.

47. Percie du Sert N, Hurst V, Ahluwalia A, Alam S, Avey MT, Baker M, et al. The ARRIVE guidelines 2.0: Updated guidelines for reporting animal research. PLoS Biol 2020;18:e3000410.

